# Multi-ancestry fine-mapping accounting for ancestral and environmental heterogeneity improves resolution and interpretation

**DOI:** 10.1101/2025.11.26.690727

**Authors:** Siru Wang, Oyesola O. Ojewunmi, Fraser J. Pirie, Ayesha A. Motala, Michele Ramsay, Andrew P. Morris, Segun Fatumo, Tinashe Chikowore, Jennifer L. Asimit

## Abstract

Amongst genome-wide association studies (GWAS) across diverse populations, allelic effect heterogeneity may arise due to differences in genetic ancestry and environmental exposures. This heterogeneity impacts the refinement of sets of potential causal variants underlying genetic associations through statistical fine-mapping. We introduce two multi-ancestry fine-mapping methods, MR-MEGAfm and env-MR-MEGAfm, allowing for multiple causal variants in a genomic region. Both methods integrate GWAS summary statistics and differing linkage disequilibrium from multiple cohorts; env-MR-MEGAfm additionally incorporates summary-level environmental covariates. Through simulations, we show that, when allelic heterogeneity is correlated with environmental exposures and ancestry, env-MR-MEGAfm yields improved resolution over MR-MEGAfm and similar resolution to SuSiEx. In twelve sex-stratified African GWAS of low-density lipoprotein cholesterol in 19,589 individuals, MR-MEGAfm and env-MR-MEGAFM (adjusting for urban status) identify five variants with posterior probability > 0.5 within two loci. One variant showed heterogeneity only due to ancestry, while three showed heterogeneity due only to urban status.

## Introduction

Genome-wide association studies (GWAS) have been successful in identifying genetic variants that are associated with many complex diseases and disease-related traits^1–4^. Many genetic variants detected as having a trait/disease-association do not play a causal role but instead are correlated with a causal variant due to linkage disequilibrium (LD). Therefore, statistical fine-mapping methods are used to construct credible sets of potential causal variants for follow-up in downstream experimental functional studies^5^.

Current fine-mapping methods can be broadly classified into single-ancestry^6–10^ and multi-ancestry fine-mapping methods^11–15^. As more diverse populations are included, increasing evidence demonstrates that multi-ancestry fine-mapping methods, which account for diverse LD patterns across ancestries, enable more powerful and accurate fine-mapping^16–18^. Multi-ancestry fine-mapping methods that allow for multiple causal variants can be grouped into two categories. The first category applies single-population fine-mapping methods to meta-analyse GWAS summary statistics using a combined ancestry LD; however, these methods do not account for allelic heterogeneity and differing LD patterns across populations^19,20^. The second category comprises Bayesian methods that integrate multiple LD structures^11,12^. Although these Bayesian-based fine-mapping methods differ in their modelling assumptions for handling the heterogeneity in allelic effects across populations, they assume that causal variants are shared across all populations^11,12^. In reality, it has been demonstrated that half of causal variants are not shared across all populations^21^. Therefore, this strong assumption potentially causes bias towards detecting shared causal variants instead of ancestry-specific causal variants. Moreover, a major limitation among these Bayesian methods is their poor computational scalability when applied to a larger number of populations.

Recently, several methods have been developed under a more realistic assumption that both shared and ancestry-specific causal variants are present across all populations. For example, MESuSiE^14^ allows for multiple causal variants present in a genomic region, including ancestry-specific causal variants, and incorporates the LD structures for each of the populations. However, MESuSiE has a key limitation: it only allows for analysis of the variants present in all populations. Additionally, MESuSiE is not scalable to a large number of variants in the region and is impractical for more than five populations. In contrast, both MGflashfm^13^ and SuSiEx^15^ overcome the restriction that variants are present across all populations. Specifically, MGflashfm and SuSiEx not only prioritise causal variants shared amongst all or a subset of ancestry groups, but also accommodate variants (non-causal or causal) that may be missing in some cohorts. However, MGflashfm is currently limited to six genetic ancestry groups, and meta-analysis across genetically similar groups is recommended prior to use.

Current multi-ancestry fine-mapping methods can identify multiple causal variants, but these methods do not account for the expected structure of heterogeneity - that is, genetically similar populations should have similar allelic effects, and populations with similar environmental exposures are expected to exhibit similar allelic effects. Furthermore, although these fine-mapping methods primarily focus on the inference of a “causal configuration”, they do not further quantify the extent of the genetic effect heterogeneity due to ancestry or environmental exposures which may interact with genetic variants^22^. Several studies have shown that some genetic loci exhibit different effects on the associated trait in populations, driven by environment or lifestyle exposure status^23,24^. Meta-analysis approaches MR-MEGA^25^ and MANTRA^26^ account for allelic heterogeneity and can be used not only for performing genetic effect heterogeneity tests, but also for fine-mapping across multiple populations, under the assumption of a single causal variant. Recently, MR-MEGA has been extended to account for not only the heterogeneity in allelic effects that are correlated with genetic ancestry but also the heterogeneity in varied environmental factors between populations - environment-adjusted MR-MEGA^27^, env-MR-MEGA^28^.

Here, we introduce powerful and scalable multi-population fine-mapping methods, MR-MEGAfm/env-MR-MEGAfm. MR-MEGAfm accounts for allelic heterogeneity that is correlated with genetic ancestry, while env-MR-MEGAfm accounts for both genetic ancestry and environmental exposures. Importantly, they allow for multiple causal variants and do not require variants to be present in all populations. Employing a stepwise selection procedure, MR-MEGAfm and env-MR-MEGAfm utilise MR-MEGA and env-MR-MEGA to perform variant selection based on an approximate conditional analysis (similar to GCTA-COJO^29^) that utilises cohort-level LD (if available). In extensive simulations, we assessed prioritisation, power and false discovery rate (FDR) of the proposed fine-mapping methods. Finally, we apply MR-MEGAfm/env-MR-MEGAfm and SuSiEx to two loci robustly associated with low-density lipoprotein (LDL) cholesterol, using GWAS summary statistics from twelve sex-stratified African cohorts consisting of a total of ∼19,000 individuals, and compare their 99% credible sets (CS99) in terms of size (number of variants) and marginal posterior probability (MPP) of causal association for variants in the CS99.

## Results

### Meta-regression fine-mapping frameworks

Assuming that a locus contains variants associated with a trait at genome-wide significance (p-value < 5 × 10^−8^), as identified using (env-)MR-MEGA^25,27^, we defined a 1Mb region centred on the lead SNP for subsequent fine-mapping. The (env-)MR-MEGAfm method is designed for fine-mapping genetic associations within a region across multiple cohorts by integrating cohort-specific GWAS summary statistics with their corresponding LD, allowing for multiple causal variants. It integrates the approximate conditional analysis approaches of GCTA-COJO^29^, only requiring GWAS summary statistics, with the meta-regression model of MR-MEGA^25^ to account for ancestral heterogeneity, or with env-MR-MEGA^27^,to account for both ancestral and environmental heterogeneity. Specifically, in the MR-MEGA framework, the allelic effects are modelled as a function of axes of genetic ancestry, derived from a matrix of mean pairwise allele frequency differences between cohorts. In the env-MR-MEGA model, the cohort-level environmental impacts, obtained by taking the mean or proportion of the individual-level environmental data within each cohort, are also included as covariates in addition to the axes of genetic ancestry. Building on this framework, (env-)MR-MEGAfm is developed using a stepwise selection approach to partition independent association signals iteratively. Additionally, by incorporating the (env-)MR-MEGA meta-regression model, (env-)MR-MEGAfm not only naturally allows assessment of variants (causal or non-causal) that are missing in some cohorts but also effectively identifies causal variants that are either shared or specific to different ancestry groups.

Specifically, for a locus containing a genome-wide significant variant, similar to the original GCTA-COJO approach^29^, (env-)MR-MEGAfm, initialises a multi-variant model by identifying the lead SNP with the smallest (env-)MR-MEGA p-value from the meta-analysis of all cohorts. Then, conditioning on the previously selected SNPs, the effect size of each SNP is estimated within each cohort by using the cohort-specific LD and GCTA-COJO^29^. Based on the conditional allelic effect estimates and their corresponding standard errors, (env-)MR-MEGA meta-analysis is then employed to select the next associated SNP (having squared multiple correlation r^2^>0.9 with previously selected SNPs) to enter the model. This procedure is iteratively repeated until no SNP attains genome-wide significance (Figure 1, Step 2.1-2.2).

**Figure 1.**
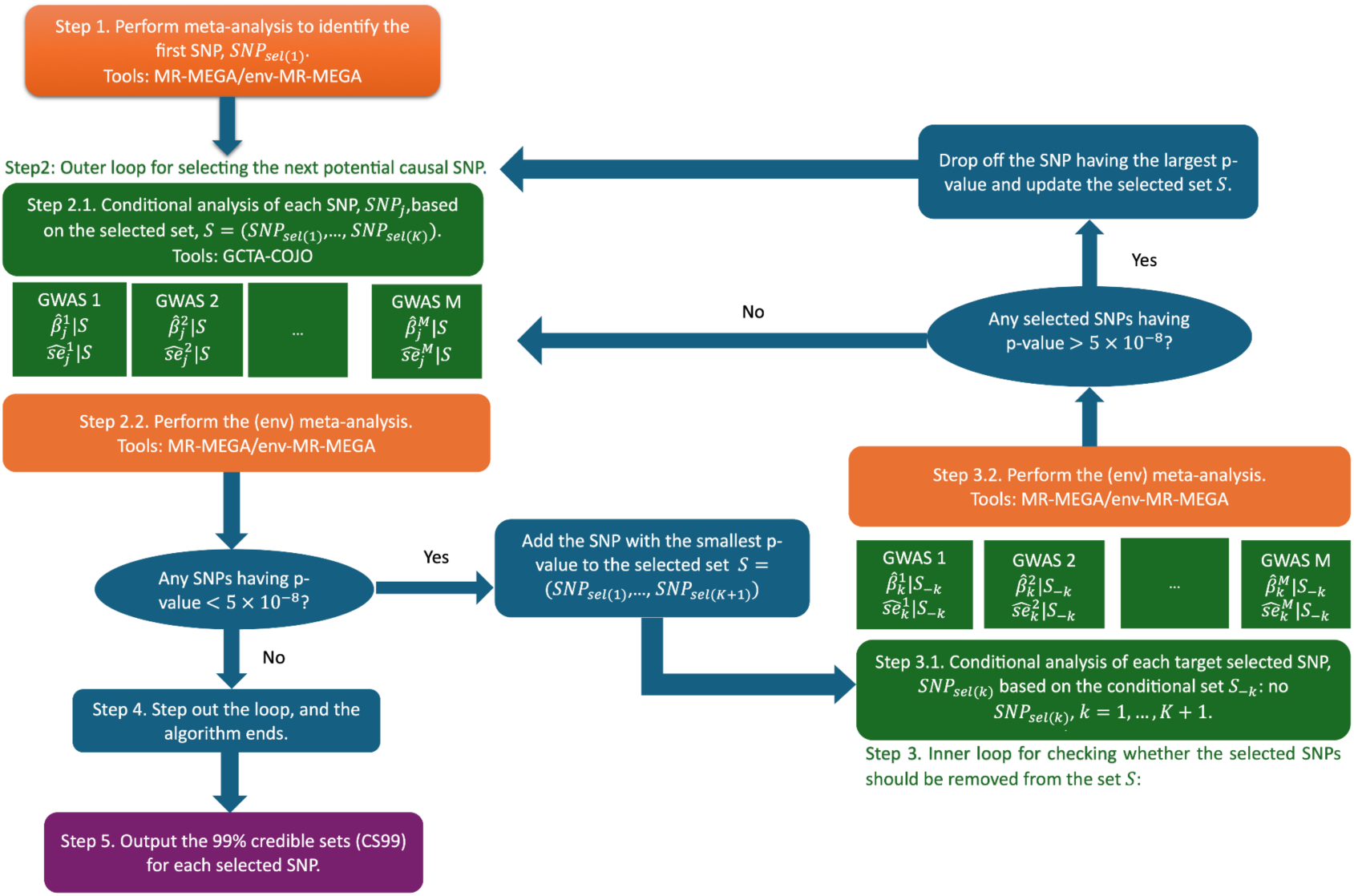
Schematic diagram of the (env-)MR-MEGA fine-mapping method.

Before continuing the variant selection procedure for the model, we assess whether any of the selected SNPs could be removed from the current set of selected SNPs. During the re-evaluation procedure, we iteratively choose one SNP from the set of currently selected SNPs, *S*, as the target SNP and the remaining selected SNPs as the conditional set, *S*_−*k*_. We estimate effect size and the corresponding standard error of the target variants, conditional on *S*_−*k*_, and assess conditional significance using (env-)MR-MEGA. If any SNP has a conditional (env-)MR-MEGA p-value > 5 × 10^−8^, the SNP with the largest p-value is removed. Otherwise, the process proceeds to the next selection step (Figure 1, Step 3.1-3.2, 4). After the iterative selection and re-evaluation procedure, we construct a 99% credible set (CS99) of each selected variant, and output the marginal posterior probabilities (MPP) for variants included in CS99, conditioned on the remaining selected variants and obtained through the re-evaluation procedure. Details on the MPP calculation and CS99 construction are provided in the second subsection of the Method and Material section, titled “The (env-) MR-MEGAfm multi-group fine-mapping”.

### Overview of simulation

To assess the performance of MR-MEGAfm and env-MR-MEGAfm, we conducted a series of simulations in a genomic region harbouring *APOE* (chr19: 44861224-45861224), and simulated genotypes of five continental African populations based on 1000 Genomes Phase 3 reference panels^30^ using Hapgen2^31^. The five populations were grouped into three clusters: the East Africa group, including Luhya in Webuye, Kenya (LWK); the West-central Africa group, including Esan in Nigeria (ESN) and Yoruba in Ibadan, Nigeria (YRI); and the West Africa group, including Gambian in Western Division, The Gambia (GWD) and Mende in Sierra Leone (MSL) (Supplementary Figure 1). The population-level LD matrices were then calculated and used as inputs to the fine-mapping methods. Using true population-matched LD in all simulations is consistent with GCTA-COJO^32^ recommendations, which suggests that the LD reference sample size is greater than 4,000 and preferably using the GWAS sample itself if the summary data come from a single cohort based GWAS. Across all populations, two causal variants were randomly selected under a unified setting with MAF > 0.05 and pairwise correlation *r^2^* < *0*.*6* within each population group in each simulation replication. Within each replication we select a sample of size 10,000 from each population and randomly split the genotype data into male and female cohorts (size 5,000 each), then generate traits within each sex-stratified cohort, either with varying effect sizes of causal variants, ranging from 0.1 to 0.3, or under different heterogeneity scenarios. Each simulation setting was replicated 300 times. Simulation design details are available in the Method and Material section.

Through extensive simulations, we assigned causal variants and varying environmental exposures to different African population groups and then compared the performance of the two proposed methods, env-MR-MEGAfm and MR-MEGAfm, with SuSiEx multi-ancestry fine-mapping^15^. SuSiEx^15^ was developed based on the single-population fine-mapping SuSiE^33^ model and allows causal variants to have varying effect sizes, including null effects, across all population groups. This design enables the method to be applied in the situation where allelic heterogeneity is impacted by ancestry or environmental exposures. We focused on comparisons with SuSiEx because other multi-ancestry fine-mapping methods exclude variants that are not present in all cohorts or are not scalable to more than a handful of populations.

For (env-)MR-MGAfm, in addition to cohort-specific GWAS summary statistics and the corresponding LD, the inputs include the axes of genetic variation and cohort-level environmental exposures. The axes of genetic variation must be derived based on the genome-wide variants with MAF>0.05 across all cohorts, rather than the variants within the specified fine-mapping region. Each fine-mapping method outputs the selected variants and their corresponding CS99. Based on these results of (env-)MR-MEGAfm and SuSiEx, we assessed the coverage, resolution, prioritisation, power, and false discovery rate (FDR). Specifically, a CS99 for each selected variant is constructed by including variants in decreasing order of their MPP until the cumulative sum first passes above 0.99. For ease in comparisons, within each replication, all CS99 are merged into the union of CS99. The coverage is estimated by the proportion of simulation replications in which the union of CS99 contains both causal variants and resolution is measured as the number of variants in the CS99 union. The prioritisation of a variant is evaluated by determining if it has evidence supporting a causal association at a pre-specified threshold of marginal posterior probability (MPP) and power is defined as the probability that a causal variant is prioritised. Similarly, FDR is defined as the probability that a prioritized variant is non-causal. Detailed definitions of these evaluation metrics are provided in the Simulation design section.

### All fine-mapping methods showed comparable resolution and calibration in the absence of heterogeneity of allelic effects due to ancestry or environment

We first examined the performance of env-MR-MEGAfm, MR-MEGAfm, and SuSiEx^15^ in a “homogeneous” setting where causal effects were the same across all African populations and were not impacted by any environmental exposures. We assess the calibration of multi-ancestry methods through simulations to estimate the coverage - the probability that a CS99 captures both causal variants. For a well-calibrated method, the expected coverage of CS99 is 0.99. Both env-MR-MEGAfm and MR-MEGAfm were well-calibrated, and both methods shared identical coverage when allelic effects of both causal variants were not correlated with the environment (Supplementary Figure 2a). Additionally, SuSiEx showed a slightly lower coverage compared to env-MR-MEGAfm and MR-MEGAfm (Supplementary Figure 2a).

Next, we examined the FDR, defined as the mean proportion of variants having marginal posterior probability (MPP) above a certain threshold (e.g. 0.5, 0.9), and power, defined as the mean proportion of causal variants having MPP above a certain threshold (e.g. 0.5, 0.9). Due to the absence of heterogeneity in allelic effects that is correlated with environment or ancestry, both env-MR-MEGAfm and MR-MEGAfm showed nearly equivalent FDR and power (Figure 2). In contrast, regardless of MPP threshold, SuSiEx had slightly inflated FDR and comparable power compared to env-MR-MEGAfm and MR-MEGAfm. The inflated FDR observed for SuSiEx is in agreement with previous results for the setting where effect sizes are highly concordant across studies^15^. To further evaluate prioritisation of causal variants, we assessed the distribution of minimum MPP across both causal variants. Under the homogeneous scenario, where heterogeneity of allelic effects was uncorrelated with either environment or ancestry, (env-)MR-MEGAfm and SuSiEx consistently produced nearly identical MPP over 300 replications (Supplementary Figure 2b) indicating no significant difference in prioritisation of both causal variants. Finally, we measured resolution by counting the number of variants in the CS99 union, obtained by merging all CS99 across selected variants in each replication. The sizes of the CS99 union from env-MR-MEGAfm were smaller than those from SuSiEx (Supplementary Figure 3b), whereas env-MR-MEGAfm had comparable resolution with MR-MEGAfm due to homogeneity of allelic effects across all population groups (Supplementary Figure 3a).

**Figure 2.**
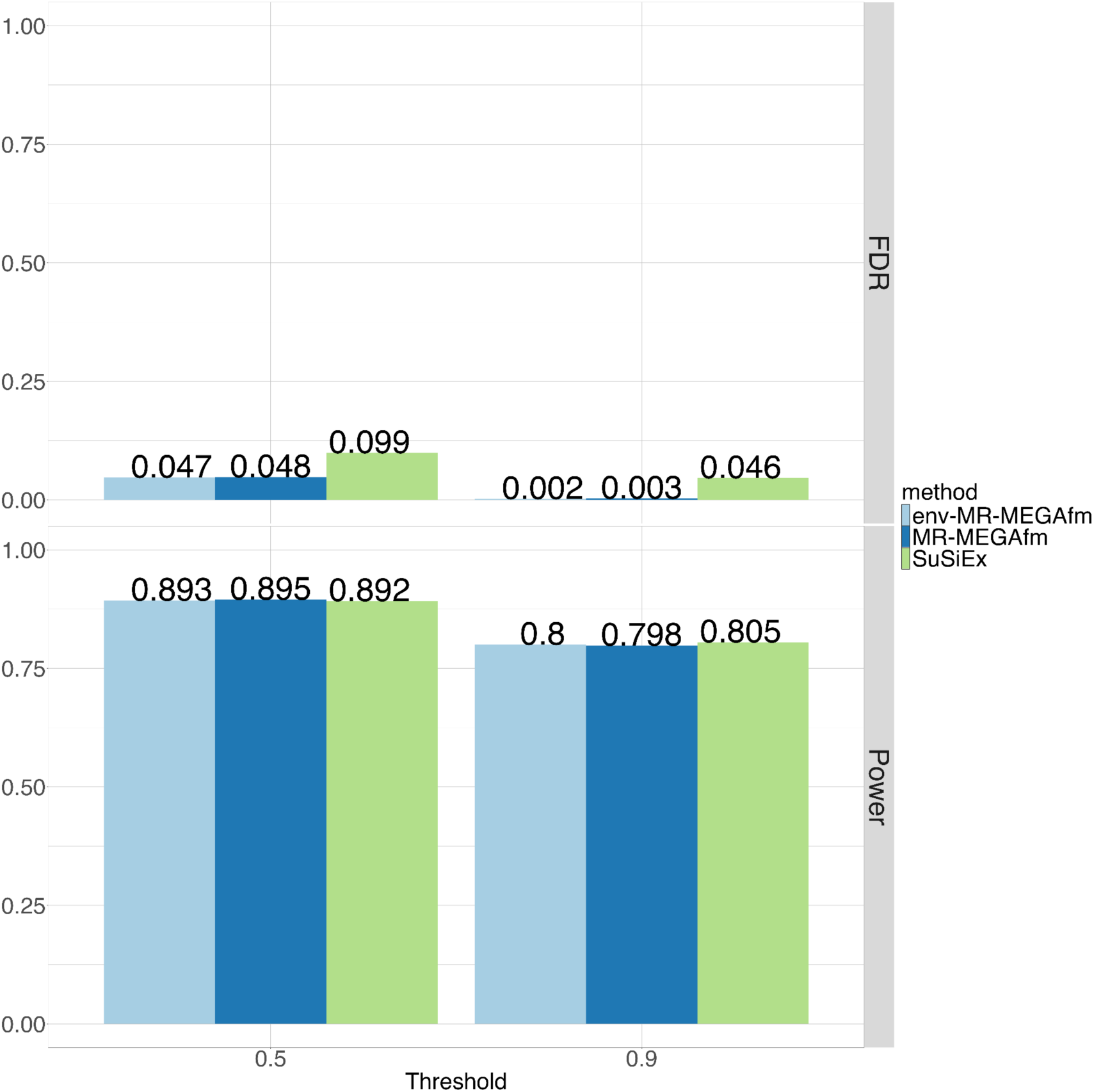
Under the homogeneity scenario where the heterogeneity of allelic effects was not correlated with either environment and ancestry, both env-MR-MEGAfm and MR-MEGAfm showed lower FDR than SuSiEx, while both methods had comparable power with SuSiEx. FDR is defined as the mean proportion of variants having marginal posterior probability (MPP) above a certain threshold that are non-causal variants, and power is measured as the mean proportion of causal variants having marginal posterior probability (MPP) above a certain threshold. FDR and power are based on 300 replications.

### Both env-MR-MEGAfm and SuSiEx outperformed MR-MEGAfm across different ancestral heterogeneity scenarios where allelic heterogeneity is correlated with varied smoking effects

To investigate the properties of MR-MEGAfm and env-MR-MEGAfm under the ancestrally homogeneous scenario where allelic heterogeneity is correlated with environment only, we treated smoking status as the environmental exposure and assigned different smoking proportions into the ten sex-stratified African cohorts, where female cohorts always had lower smoking proportions than male cohorts (Supplementary Table 1). Here, we still compare the performances between our proposed fine-mapping methods, (env-)MR-MEGAfm, and SuSiEx in the series of simulation settings where allelic heterogeneity is impacted by smoking effects.

Regardless of causal effect sizes, both env-MR-MEGAfm and MR-MEGAfm were consistently well-calibrated for CS99, with a coverage greater than 0.98 (Supplementary Table 2a). Specifically, the upper 95% confidence limit for coverage from both methods included 100%. Additionally, SuSiEx always showed comparable calibration performance with the proposed methods (Supplementary Table 2a). Furthermore, we evaluated the FDR and power of the three fine-mapping methods, using a moderate threshold (0.5) and a common stringent threshold (0.9) (Supplementary Table 3). When allelic heterogeneity is correlated with environment (smoking effects) only, env-MR-MEGAfm had higher power than MR-MEGAfm while it shared comparable power with SuSiEx. Specifically, as causal allelic effects range from 0.1 to 0.3, the three methods showed a notable increase in power regardless of thresholds. However, there was no significant difference in FDR between these three methods regardless of the causal allelic effects. As heterogeneity in allelic effects was correlated with smoking effects, env-MR-MEGAfm demonstrated higher gains in resolution and prioritisation over MR-MEGAfm (Supplementary Figure 4a, Supplementary Figure 4b). Additionally, across varied causal allelic effect settings, SuSiEx showed comparable resolution performance (Supplementary Figure 4a, Supplementary Figure 5), and prioritisation performance to env-MR-MEGAfm (Supplementary Figure 4b). As expected, the increasing causal effect sizes led to a significant reduction in the sizes of the CS99 and more noticeable improvement in the prioritisation of both causal variants for the three fine-mapping methods.

To further evaluate the performance of proposed fine-mapping methods under the scenarios where both smoking effects and ancestry have effects on the allelic heterogeneity, we introduced two heterogeneity scenarios: west-central African and non-ancestral African scenarios (Supplementary Table 4). For example, in the west-central African scenario, sub-populations of smokers from west-central Africa (ESN and YRI) have a trait associated with the specific causal variants, while all other populations show no association at these variants. For each heterogeneity scenario, we consider the allelic effect heterogeneity due to smoking effects, in addition to ancestral effects. All quantitative traits under different heterogeneity scenarios were simulated with the causal effect size of 0.3.

Across all heterogeneity scenarios where heterogeneity of allelic effects is impacted by both ancestry and smoking effects, all three fine-mapping methods consistently had well-calibrated coverage at 99% (Figure 3a, Supplementary Table 2b). The env-MR-MEGAfm and SuSiEx showed nearly equivalent coverage, with slight gains over MR-MEGAfm under each heterogeneity scenario.

**Figure 3.**
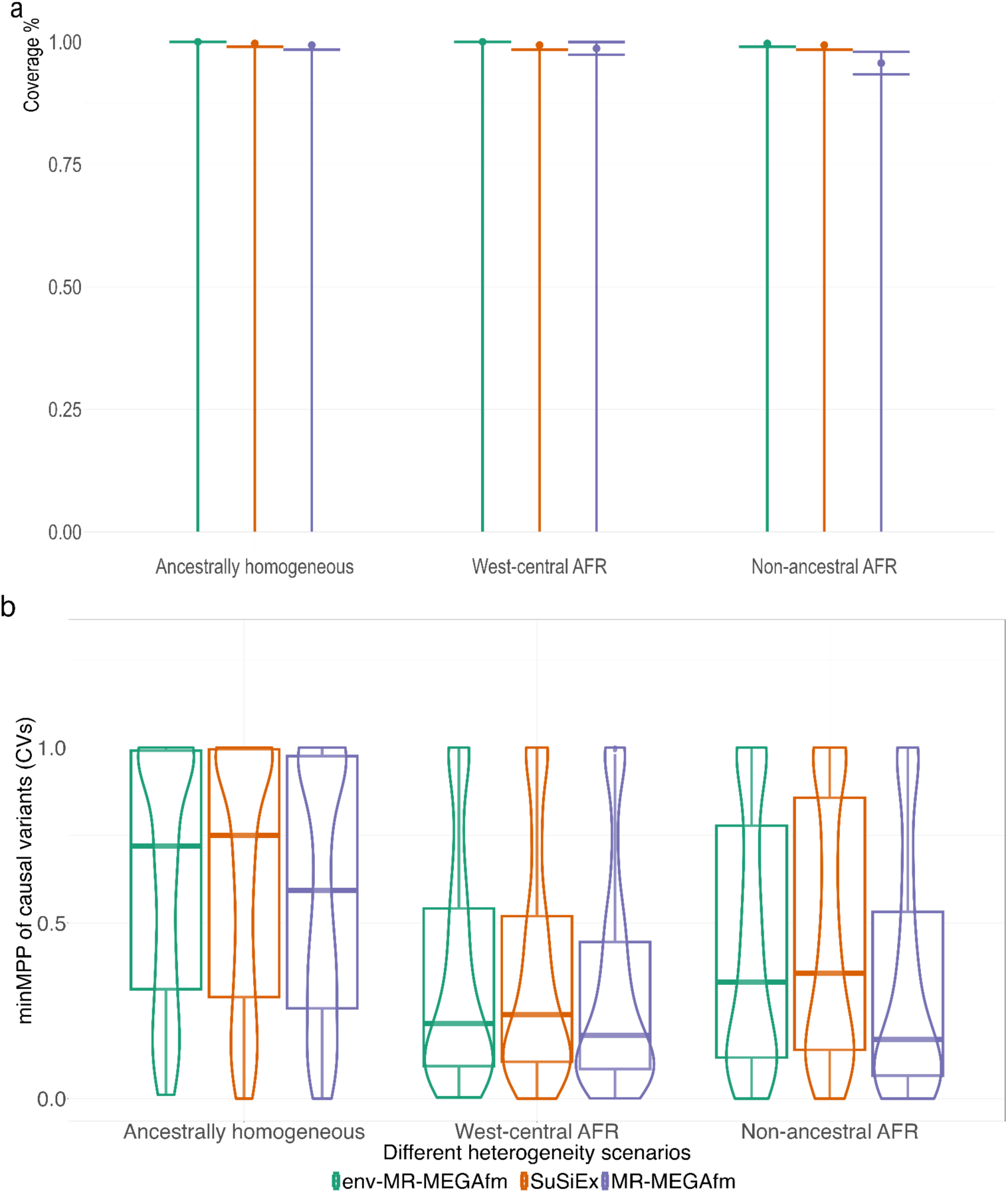
Across different heterogeneity scenarios in which smoking status had an impact on allelic heterogeneity, env-MR-MEGAfm, MR-MEGAfm and SuSiEx were well-calibrated. Additionally, env-MR-MEGAfm consistently had higher gains in prioritisation than MR-MEGAfM. a. Coverage from different heterogeneity scenarios. Coverage is measured as the probability that both causal variants are captured by CS99 union estimated over 300 replications. Data are presented as the proportion of replications in which the CS99 union contains all causal variants±SEM, where SEM is the standard proportion error bound of a 95% confidence interval based on 300 replications. **b Prioritisation Assessment.** The boxplots show MPP distribution of the three fine-mapping methods, and the median is given by the centre line, upper and lower quartiles are the box limits, and whiskers are at most interquartile range. This suggests that env-MR-MEGAfm and SuSiEx perform comparably under the ancestrally homogeneous and west-central Africa scenarios, and both outperform MR-MEGAfm at prioritising both causal variants.

Under each heterogeneity scenario, we evaluated the results of env-MR-MEGAfm, MR-MEGAfm and SuSiEx in terms of FDR and power (Table 1), and, as expected, env-MR-MEGAfm and SuSiEx consistently produced lower FDR and higher power than MR-MEGAfm regardless of the threshold. Additionally, as the ancestrally homogeneous scenario included more African populations with genetic associations, these fine-mapping methods demonstrated slightly increased power and decreased FDR. Compared to the ancestrally homogeneous scenario, the west-central Africa scenario included more African populations with null genetic associations with a trait. This increases the challenge in multi-ancestry fine-mapping to identify the potential causal variants, resulting in substantially reduced power under the ancestry-specific heterogeneity scenario (west-central Africa scenario). To mimic the real sample sizes across different African populations from east Africa, south Africa and west Africa, we set two sample size settings in which each sex-stratified cohort contains smaller individual samples ranging from 1,000 to 3,000 under the west-central Africa scenario (Supplementary Table 5). Compared to the west-central Africa scenario with sample size of 5,000, although all fine-mapping methods were robust to FDR, they had minor reductions in coverage and power, especially for power under the stringent threshold (0.9) (Supplementary Table 6).

**Table 1.** Under each heterogeneity scenario where smoking status was included as an environmental exposure, env-MR-MEGAfm provided slightly higher power and lower FDR compared to MR-MEGAfm, as expected. However, SuSiEx consistently had nearly equivalent FDR and power with env-MR-MEGAfm, regardless of thresholds. FDR is defined as the mean proportion of variants having marginal posterior probability (MPP) above a certain threshold that are non-causal variants, and power is measured as the mean proportion of causal variants having MPP above a certain threshold. FDR and power are based on 300 replications.

| Scenarios | Ancestrally homogeneous |  | West-central Africa |  | Non-ancestral Africa |  |
| --- | --- | --- | --- | --- | --- | --- |
| Threshold (0.5) | FDR | Power | FDR | Power | FDR | Power |
| MR-MEGAfm | 0.088 | 0.740 | 0.132 | 0.485 | 0.165 | 0.535 |
| env-MR-MEGAfm | 0.072 | 0.795 | 0.128 | 0.530 | 0.102 | 0.630 |
| SuSiEx | 0.082 | 0.800 | 0.129 | 0.527 | 0.092 | 0.635 |
| Threshold (0.9) |  |  |  |  |  |  |
| MR-MEGAfm | 0.010 | 0.583 | 0.011 | 0.330 | 0.030 | 0.403 |
| env-MR-MEGAfm | 0.003 | 0.635 | 0.013 | 0.365 | 0.006 | 0.478 |
| SuSiEx | 0.018 | 0.647 | 0.021 | 0.378 | 0.006 | 0.495 |

Across different heterogeneity scenarios, env-MR-MEGAfm tended to produce smaller credible sets than MR-MEGAfm, and there were only slight differences between env-MR-MEGAfm and SuSiEx (Supplementary Figure 6). Furthermore, as causal variants exhibited genetic associations in more African populations (e.g. west-central Africa and ancestrally homogeneous scenarios), CS99 union sizes yielded by these fine-mapping methods became notably smaller.

As expected, across all heterogeneity scenarios where smoking effects (the environmental exposure) had an impact on heterogeneity of allelic effects, env-MR-MEGAfm and SuSiEx produced improved prioritisation of both causal variants compared to MR-MEGAfm (Figure 3b). Additionally, under each heterogeneity scenario, env-MR-MEGAfm and SuSiEx produced comparable MPP of causal variants across 300 replications. More specifically, env-MR-MEGAfm showed slight gains in prioritisation of causal variants compared to SuSiEx. (Supplementary Figure7).

### The cohort-matched LD utilized in all fine-mapping methods outperformed the approximate LD

After examining the calibration of MR-MEGAfm and env-MR-MEGAfm by varying causal effects and heterogeneity scenarios, we confirmed that the proposed fine-mapping methods consistently were well-calibrated when the true cohort-level LD patterns were employed. However, individual-level genotype data for cohorts may not be available, and reference LD or combined LD may be considered to reduce the computational burden of multiple LD matrices. We compare the performance of the proposed fine-mapping methods with the combined LD as a replacement for the cohort-matched LD under the ancestrally homogeneous scenario where allelic effect heterogeneity is correlated with smoking effects only. Under this setting, env-MR-MEGAfm and SuSiEx outperformed MR-MEGAfm. However, as expected, using the combined LD in all fine-mapping methods resulted in substantial losses in resolution, prioritization, coverage and power compared with the true cohort-level LD. (Supplementary Table 7, Supplementary Figure 8a, Supplementary Figure 8b). Therefore, we recommend the use of true cohort-level LD in fine-mapping.

### Fine-mapping LDL cholesterol genetic associations accounting for urban status

We applied MR-MEGAfm, env-MR-MEGAfm, and SuSiEx^15^ to fine-map putative causal variants for the LDL-cholesterol trait. Specifically, we used summary statistics from twelve sex-stratified GWAS of LDL-cholesterol, consisting of 19,589 participants, from East, West and South African cohorts from the Africa Wits-INDEPTH partnership for Genomics studies (AWI-Gen) consortium^34,35^, South African Zulu cohorts: the Durban Diabetes Case Control Study (DCC) and the Durban Diabetes Study (DDS)^36^ and Uganda: Uganda Genome Resource^37–39^. The axes of genetic variation further demonstrate these cohorts were grouped into three main clusters: East Africa (AWI-Gen East, Uganda), West Africa (AWI-Gen West) and South Africa (AWI-Gen South, Zulu Case and Control) (Figure 4a). We focused on the two genomic regions centred on the two genome-wide significant loci on chromosome 1 (*PCSK9*: rs28362286) and chromosome 19 (*APOE:* rs7412), where the two loci identified by env-MR-MEGA^27^ meta-analysis showed significant evidence of allelic heterogeneity driven by urban status, the proportion of study participants who live in an urban area. For env-MR-MEGAfm, we used urban status as an environmental variable within the specific genomic regions across the twelve sex-stratified African populations. Additionally, we utilised SABER^40^ to identify the genomic region boundaries for the two loci and limited fine-mapping to 1Mb regions centred on the two loci.

**Figure 4.**
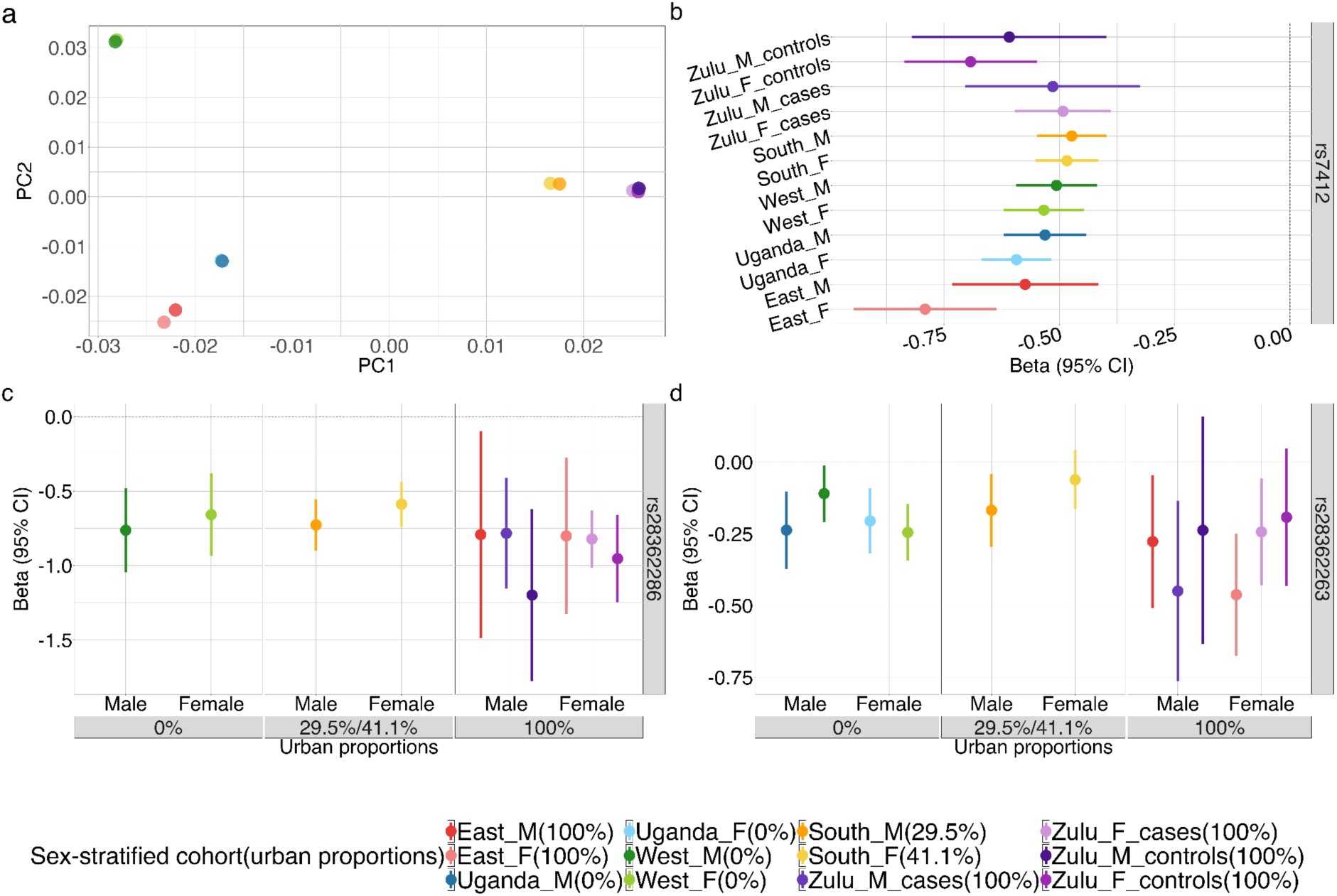
Axes of genetic variation and allelic heterogeneity due to urban status across twelve sex-stratified African cohorts. **a.** Axes of genetic variation: axes of genetic variation showing separation into three distinct clusters representing cohorts from West Africa (West_M &West_F), East Africa (East_M & East F and Uganda_M & Uganda_F), and south Africa (South_M & South_F, controls for Zulu_M & Zulu_F and cases for Zulu_M & Zulu_F). **b.** Allelic heterogeneity across all cohorts exhibiting the negative effects for rs7412 with LDL-cholesterol trait. **c & d.** Allelic heterogeneity due to urban status with negative effects for rs28362286 and rs28362263 is more apparent towards urban residents’ cohorts. The X-axis represents the urban proportions, while the Y-axis represents allelic effect sizes and their 95% credible intervals (CI) across the twelve cohorts. For **b**, **c** and **d,** 95% CIare provided as error bars for each effect estimate. Source data for **b**, **c** and **d** are available in Supplementary Table 8. Source data in Supplementary Data 1.2.

In the *PCSK9* region, all fine-mapping methods prioritised rs28362286 (MPP=1). In addition, env-MR-MEGAfm and SuSiEx gave high confidence for rs28362263 (Table 2). These two variants have been previously prioritised for LDL-cholesterol in African ancestries by MGflashfm^13^. Specifically, in fine-mapping LDL in *PCSK9* and *USP24*, the prioritised rs28362263, a missense variant, is only polymorphic in Africa (AFR) and Hispanic/Latino (HIS) (pooled MAF =0.050). Stop-gain variant rs28362286 is also only polymorphic in AFR (MAF=0.00798) and was also prioritised by the Global Lipids Genetics Consortium (GLGC) African ancestry analysis^41^, but not by the GLGC multi-ancestry analysis. For the *APOE* region, the three methods prioritised rs7412 and rs429358 for a causal association with LDL cholesterol (MPP=1), which are already established causal variants and define *APOE ε*-alleles^42–45^ (Table 2, Supplementary Data 1.1). SuSiEx^15^ prioritised an additional variant on chromosome 1 with the corresponding position 55561346 (MPP = 0.826), and the variant only exists in the West Africa population group. MR-MEGAfm and env-MR-MEGAfm cannot detect this variant due to the limited number of cohorts in which the variant is present. Based on the identified variants, we constructed the corresponding credible sets and listed all variants having MPP>0.5 (Table 2). For these variants, the corresponding conditional p-values of association attain genome-wide significance.

**Table 2.** Fine-mapping of credible set SNPs with MR-MEGAfm/env-MR-MEGAfm and SuSiEx in LDL-cholesterol summary statistics of African individuals.

|  | The number of cohort that SNP are present | Posterior inclusion probability (PIP) | MPP | Allelic heterogeneity due to ancestry p-values | MPP | Allelic heterogeneity due to ancestry p-values | Allelic heterogeneity due to urban status p-values |
| --- | --- | --- | --- | --- | --- | --- | --- |
| rsID |  | SuSiEx | MR-MEGAfM |  | env-MR-MEGAfM |  |  |
| PCSK9 | CHR1:55029215-56029215 |  |  |  |  |  |  |
| rs28362286 | 10 | 1 | 1 | 0.858 | 1 | 0.470 | 0.036 |
| rs28362263 | 12 | 0.927 | 0.65 | 0.113 | 0.873 | 0.103 | 0.035 |
| 1:55561346 | 2 | 0.826<br>(Exist in west AFR) |  |  |  |  |  |
| APOE | CHR19:44912079-45912079 |  |  |  |  |  |  |
| rs7412 | 12 | 1 | 1 | 0.008 | 1 | 0.098 | 0.002 |
| rs429358 | 12 | 1 | 1 | 0.355 | 1 | 0.539 | 0.307 |
| rs769455 | 12 | 0.681 | 0.686 | 0.498 | 0.724 | 0.016 | 0.731 |

Amongst the prioritised variants having MPP>0.5, MR-MEGAfm and env-MR-MEGAfm allow evaluation of the contribution of ancestry and the environmental variable (urban proportions) to allelic effect heterogeneity using a nominal p-value threshold of 0.05. Four variants showed allelic effect heterogeneity due to ancestry or environment (Table 2). Specifically, allelic effect heterogeneity due to ancestry was detected by MR-MEGAfm at rs7412 (MPP=1 in *APOE*), whereas env-MR-MEGAfm detected allelic effect heterogeneity due to ancestry only at rs769455 (MPP=0.724 in *APOE*). Additionally, env-MR-MEGAfm showed allelic effect heterogeneity due to environment only for three variants: rs28362286, rs28362263 and rs7412. We further explored the details of the prioritised variants having significant heterogeneity of allelic effects. For rs7412, the effect sizes from four Zulu cohorts had wider confidence intervals than those from other African population groups due to small sample size. The effect sizes of rs7412 from the remaining African population groups are strongly associated with decreased LDL-cholesterol with females from East Africa having the strongest negative effect (Figure 4b, Supplementary Data 1.2). Both rs28362286 and rs28362263 across all cohorts exhibit negative allelic effects on LDL-cholesterol, with stronger negative effects observed in populations with higher urban proportions (Figure 4c-4d, Supplementary Data 1.2).

## Discussion

We have introduced multi-ancestry fine-mapping methods, MR-MEGAfm and env-MR-MEGAfm, which allow for heterogeneity in allelic effect sizes to be correlated with genetic ancestry and environmental exposures by integrating approximate conditional analyses with MR-MEGA/env-MR-MEGA models. As with many current fine-mapping methods, (env-)MR-MEGAfm methods make use of GWAS summary statistics and the associated LD patterns to approximate the conditional effect size of each genetic variant across a locus. The use of summary-level data makes these methods easily scalable to large biobank datasets. Most multi-ancestry fine-mapping methods that allow for multiple causal variants have a limitation that all variants must be present across all cohorts. This restriction becomes increasingly prohibitive as the number of studies and study diversity increases. Our methods, as well as SuSiEx^15^ and MGflashfm^13^, overcome this limitation and allow for a variant to be missing in some cohorts. Although not all variants are required to exist across all cohorts, env-MR-MEGAfm can only detect causal variants existing in at least six cohorts when accounting for one environmental covariate and two axes of genetic variation, whereas MR-MEGAfm requires causal variants to be present at least five cohorts assuming the same two axes of genetic variation. This restriction stems from the fact that (env-)MR-MEGAfm methods are built upon the MR-MEGA^25^ and env-MR-MEGA^27^ meta-regression frameworks.

Through extensive simulations incorporating varied smoking effects across different heterogeneity scenarios, env-MR-MEGAfm demonstrated significant improvements in power, FDR, resolution and prioritisation of the causal variants compared to MR-MEGAfm while producing comparable results with SuSiEx^15^. Although SuSiEx does not account for environmental effects, it estimates the posterior inclusion probability by leveraging causal probabilities across cohorts. Consequently, even when causal variants have low causal probabilities in some cohorts due to environmental impacts, the overall posterior inclusion probabilities of these causal variants produced by SuSiEx remain robustly high. Unlike SuSiEx, (env-)MR-MEGAfm detects genetic association by allowing for environmental and ancestral heterogeneity in allelic effects between population groups. Therefore, (env-)MR-MEGAfm can quantify the extent of allelic heterogeneity due to ancestry or environment. Specifically, both MR-MEGA^25^ and env-MR-MEGA^27^ demonstrated increased power to detect genetic associations when allelic effect heterogeneity is correlated with ancestry or environment. When ancestry and environment did not impact heterogeneity in allelic effects, our proposed fine-mapping methods, MR-MEGAfm/env-MR-MEGAfm, retained well-calibrated coverage. Furthermore, they shared comparable power, resolution and prioritisation of causal variants with slight improvements over SuSiEx.

In applications to GWAS of LDL cholesterol across African populations, urban status was used as an environmental exposure. Consistent with simulation results, (env-)MR-MEGAfm and SuSiEx prioritized nearly identical variants in *PCSK9* and *APOE* regions, except for one variant found only in the West African populations. The prioritized variants detected by both (env-)MR-MGAfm and SuSiEx have been previously reported as causal for LDL cholesterol^13,42,43^. Compared with SuSiEx, env-MR-MGEAfm and MR-MEGAfm additionally assessed the ancestral and/or environmental contribution to allelic heterogeneity in these variants. Specifically, in the investigation into gene-environment interactions involving LDL-cholesterol, independent loci *APOE*: rs7412, *PCSK9:* rs28362263 and rs28362286 have been shown to interact with urban status and contributed to allelic effect heterogeneity across cohorts.

In summary, (env-)MR-MEGAfm are well calibrated and improve the power of fine-mapping that integrates GWAS summary statistics from diverse populations under the assumption that heterogeneity in allelic effects is correlated with ancestry and environmental exposures. When the trait is only impacted by genetic ancestry, both MR-MEGAfm and env-MR-MEGAfm improve coverage and resolution and can detect ancestral heterogeneity; env-MR-MEGAfm can also capture environmental contributions to allelic heterogeneity. Furthermore, when the trait is affected by both environmental exposures and genetic ancestry, env-MR-MEGAfm produces the improved resolution and prioritization of the causal variants compared with MR-MEGAfm. As more GWAS from increasingly diverse populations and the associated cohort-level environmental exposures are available, (env-)MR-MEGAfm enables analysis of large and complex genomic loci and facilitates the prioritisation of functionally important causal variants that impact complex human diseases.

## Supporting information

Supplementary Daata 1.1

Supplementary Daata 1.2

## Resource availability

### Lead contact

Requests for further information and for resources should be directed to and will be fulfilled by the lead contact, Jennifer Asimit.

### Material availability

This study did not generate new unique reagents.

### Data and code availability

Our proposed (env-)MR-MEGAfm method, envMRMEGAfm^46^, is freely available as an R library at https://github.com/SiruRooney/envMRMEGAfm. The 1000 Genomes cohorts used in this paper are available from the developers of MAGMA (https://vu.data.surfsara.nl/index.php/s/ePXET6IWVTwTes4/download). Plink is freely available at https://www.cog-genomics.org/plink/. Hapgen2 is freely available at https://mathgen.stats.ox.ac.uk/genetics_software/hapgen/hapgen2.html. SuSiEx is available at https://github.com/getian107/SuSiEx.

## Acknowledgements

This research is funded by the UK Medical Research Council (MR/W02098X/1). SW and JA are also funded by the UK Medical Research Council (MC_UU_0004/1). For the purpose of Open Access, the authors have applied a CC BY public copyright licence to any Author Accepted Manuscript version arising from this submission.

## Method and material

In our stepwise selection procedure, we select SNPs to enter the model based on (env-)MR-MEGA p-values, where SNP effect estimates from each cohort are conditioned on all previously selected SNPs in the model. Within each cohort, we follow the estimation of joint and conditional SNP effects that are described in GCTA-COJO^29^ and provide details here.

### Estimating the joint multi-SNP effects and conditional single-SNP effects for a quantitative trait within each cohort

We provide estimation details here, which follow those of GCTA-COJO^29^ that was developed for a single genetic ancestry. Within a GWAS file, each SNP is independently tested for association with a trait based on a linear single-SNP model

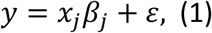

where *x_j_* = (*x_1j_*, *x_2j_*,…, *x_*n*j_*)*^T^*, *x_ij_* is the genotype score for the *j*-th SNP of the *i*-th individual; and *β_j_* is the marginal effect of the *j*-th SNP.

Considering the joint effects of multiple genetic variants on a quantitative trait, we have

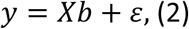

where *y* = {*y*_*i*_} is an *n* × *1* vector of phenotypes, *X* = {*x*_*ij*_} is an *n* × *J* genotype matrix and *b* = {*b_j_*} is a *J* × *1* vector of joint-SNP effects. The joint effects of multiple SNPs can be estimated by the least-squares method as

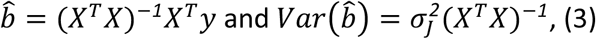

Where 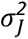 is the residual variance in the joint analysis. In a multiple regression model, 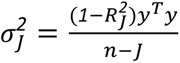 where 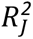 refers to the coefficient of determination of a multiple regression model and 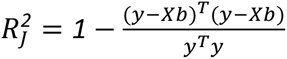. Among these *J* SNPs, the *j*-th SNP effect, conditioned on the set of the remaining (*J* − *1*) SNPs, can be expressed by

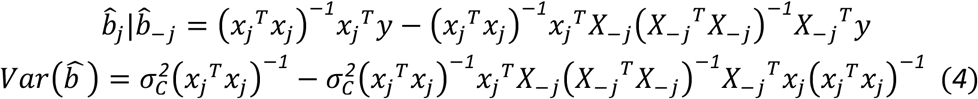

where 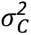 is the residual variance in the conditional analysis and *X*_−*j*_ is the genotype data of the (*J* − *1*) SNPs.

For all SNPs present in a GWAS, we can convert joint effects expressed in the equation (3) into the marginal effect of each SNP in a matrix form as

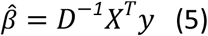

Where 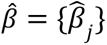 is an *J* × *1* vector of marginal SNP effects; *D* = {*D_j_*} is the diagonal matrix with 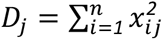. In the single-SNP analysis, the squared standard error 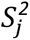 of 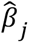 can be obtained by 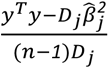 so that

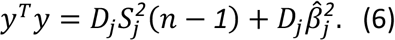

If the quantitative trait measurements are unavailable, *y*^*T*^*y* may be approximated by the median of 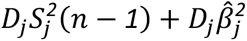 across all SNPs.

Based on equation (5), we have that the estimated joint effects of multiple SNPs *b* and the single-SNP effect conditioning on (*J* − *1*) SNPs can also be expressed by the summary statistics from a GWAS and the individual-level genotype data even without the phenotype data. The equations (3) and (4) would be transformed into

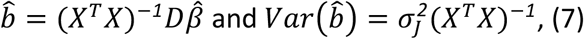

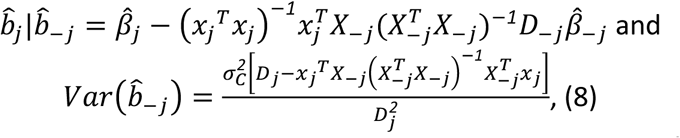

where *D*_−*j*_ is a (*J* − *1*) × (*J* − *1*) diagonal matrix without 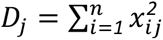 and 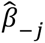 is the marginal effects of (*J* − *1*) SNPs. For the residual variance in the conditional analysis, 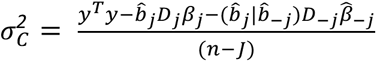.

Sometimes, we are not able to collect the pooled individual-level genotype data so that 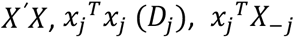 and 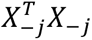 cannot be obtained. In this situation, as in GCTA-COJO^29^, we approximate information from the pooled individual-level genotype data *X* by the genotype data from the reference panel with sample size *m*, denoted by *W* = {*w_*i*j_*}. Additionally, the sample sizes of those SNPs having the pooled genotypes would be estimated as well. By equation (6), for the *j*-th SNP, 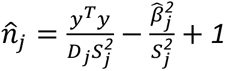. Then the variance-covariance matrices of SNP genotypes 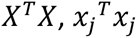 and 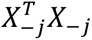 would be approximated by 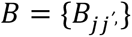, where 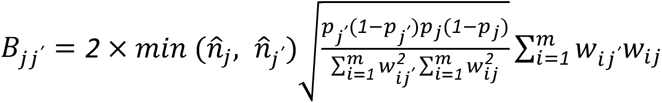 with *p_j_* and 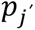 being the allele frequencies of the *j*-th SNP and the *j^ʹ^*-th SNP, respectively. In the conditional analysis, 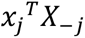 would be approximated by 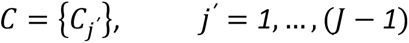, where 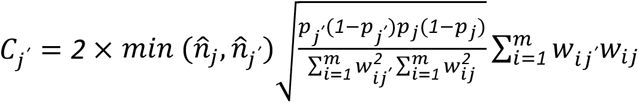.

### The (env-) MR-MEGAfm multi-group fine-mapping

To enable summary statistics-based fine-mapping across multiple cohorts that accounts for heterogeneity due to genetic ancestry or environmental exposures, we developed MR-MEGAfm and env-MR-MEGAfm, building on the MR-MEGA^25^ and env-MR-MEGA^27^ meta-regression frameworks. The (env-)MR-MEGAfm allows for variants that are not present across all cohorts, provided that the number of cohorts in which a variant is present, denoted by *M*, exceeds *T* + *S* + 2, where *T* is the number of axes of genetic variation and *S* is the number of environmental exposures. Accordingly, for MR-MEGAfm, at least five cohorts (*M* = *5*) are required when including two genetic ancestral covariates only (*T* = *2* and *S* = *0*). For env-MR-MEGAfm, at least six cohorts (*M* = *6*) are needed assuming two axes of genetic variation (*T* = *2*) and one environmental exposure (*S* = *1*). More specifically, in MR-MEGA and env-MR-MEGA frameworks, the allelic effects are modelled as a function of axes of genetic ancestry, which are derived from a matrix of mean pairwise allele frequency differences between cohorts. In the env-MR-MEGA model, the cohort-level environmental impacts are obtained by taking the mean or proportion of the individual-level environmental data within each cohort, and are included as covariates in addition to the axes of genetic ancestry.

Both MR-MEGAfm and env-MR-MEGAfm utilize a stepwise selection strategy within a multiple regression framework. In this approach, the SNP with the smallest (env-)MR-MEGA meta-analysis p-value that is also below the genome-wide significance threshold is iteratively selected from all SNPs in the region. After this variable selection stage, credible sets are constructed for each selected SNP. The detailed variable selection procedure is described below, and a schematic diagram of the (env-)MR-MEGAfm variable selection stage is provided in the third subsection of the Method and Material section, titled “ (env-)MR-MEGAfm variable selection algorithm details”.

(1) Start with a model which includes the most significant SNP in the region, identified by its smallest p-value below a genome-wide significance threshold of *5* × *10*^−*8*^ in the (env-)MR-MEGA meta-analysis. This SNP would be treated as the first selected SNP. Subsequently, the remaining SNPs within the region are scanned using the following steps.
(2) Step into the outer loop, repeatedly selecting the next potential SNP while conditioning on the previously selected SNPs.

(2.1) In the *k*-th iteration, calculate the allelic effects, and their squared standard errors for the remaining SNPs, conditioned on the set of previously selected SNPs, *S* = (*SNP*_*sel*(_*_1_*_)_,…, *SNP*_*sel*(*k*−_*_1_*_)_), within each cohort, as described in the “Estimating the joint multi-SNP effects and conditional single-SNP effect for a quantitative trait within each cohort” section. To avoid collinearity, across *M* cohorts, if the squared multiple correlation *r^2^* between a target SNP and these selected SNPs or the absolute correlation between a target SNP and one of previously selected SNPs exceeds a cut-off value, such as 0.9, the effect and the squared standard error for the target SNP will be set to NA.
(2.2) In the framework of (env-)MR-MEGA, the allelic effects of the target *SNP_j_*, 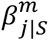, *j* = *1*,…, *J* − *k* + *1*, conditioned on (*k* − *1*) selected SNPs across *M* cohorts, *m* = *1*,…, *M*, weighted by their squared standard errors 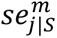, would be modelled by the linear model of (env-)MR-MEGA. MR-MEGA meta-regression model includes *T* axes of genetic variation, while env-MR-MEGA meta-regression model includes *T* axes of genetic variation and *S* environmental exposures.

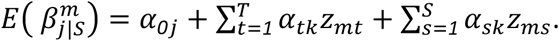

Then, select the most significant SNP with the smallest p-value from the (env-)MR-MEGA association test, provided it is below the genome-wide significance threshold of *5* × *10*^−*8*^, to enter the temporary set of the selected SNPs *S*, *S* = (*SNP*_*sel*(_*_1_*_)_,…, *SNP*_*sel*(*k*)_).
(3) Based on the temporary set of the selected SNPs (size *k*), re-evaluate all SNPs in the inner loop to identify any selected SNPs that are no longer significant and should be removed from the temporary set.

(3.1) For each cohort, iteratively select one 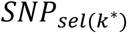 from the temporary set *S* as the target SNP. Re-calculate the allelic effect and the corresponding standard error for this SNP, conditioned on the set of the remaining selected SNPs, 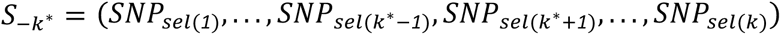, as described in step (2.1). (3.2) Across *M* cohorts, fit the 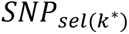 in the (env-)MR-MEGA meta-regression model, as described in step (2.2). Based on the results of the (env-)MR-MEGA meta-analysis, optimize the temporary set of the selected SNPs by removing the SNP having the largest p-value that exceeds the genome-wide significance threshold of *5* × *10*^−*8*^.
(4) Repeat steps (2) and (3) until no SNPs can be added or removed from the model.

After this variable selection stage, we turn to construct a 99% credible set (CS99) for the causal variant represented by each variant in the set of selected SNPs, *S* = (*SNP*_*sel*(_*_1_*_)_, *SNP*_*sel*(_*_2_*_)_,…, *SNP*_*sel*(*K*)_). This is done by iteratively removing one SNP, *SNP*_*sel*(*k*)_, *k* = *1*,…, *K*, from the set and considering the remaining selected SNPs as the updated set *S*_−*k*_ = (*SNP*_*sel*(_*_1_*_)_,…, *SNP*_*sel*(*k*−_*_1_*_)_, *SNP*_*sel*(*k*+_*_1_*_)_,…, *SNP*_*sel*(*K*)_). Each removed *SNP*_*sel*(*k*)_, *k* = *1*,…, *K*, represents a single causal variant, and for each *SNP_j_*, *j* = *1*,…, *J* − *K* + *1*, in the region we calculate Bayes’ factors (BF) conditional on the remaining selected SNPs in *S*_−*k*_, followed by marginal posterior probabilities (MPP) of causality, *PP_j_*|*S*_−*k*_.

Specifically, for each SNP *j* in the region, we estimate its effect and the corresponding squared standard error conditional on each updated set *S*_−*k*_ within each cohort, followed by (env-)MR-MEGA meta-analysis to calculate the conditional association test statistic *X_j_*|*S*_−*k*_. For MR-MEGA meta-analysis, the conditional BF is given by 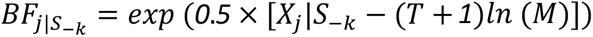, where *T* is the number of genetic variation (PCs) and *M* is the number of cohorts^25^; for env-MR-MEGA meta-analysis, the conditional BF is given by 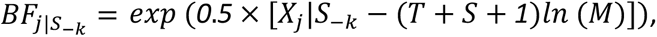, where *S* is the number of environmental exposures. After calculating the conditional Bayes factors (BFs) for all variants in the region, determine the MPP of SNP *j* using 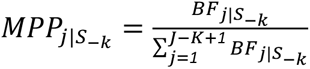, by assuming a single causal variant in the region when conditioning on the other selected variants. Finally, we construct a CS99 by setting a 99% cumulative MPP threshold (i.e. cumulative MPP=0.99) for each *S*_−*k*_and sorting the selected SNPs by decreasing 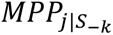. Then SNPs are selected from the sorted list until the cumulative sum of their 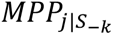 first exceeds 0.99

### The (env-)MR-MEGAfm variable selection algorithm details

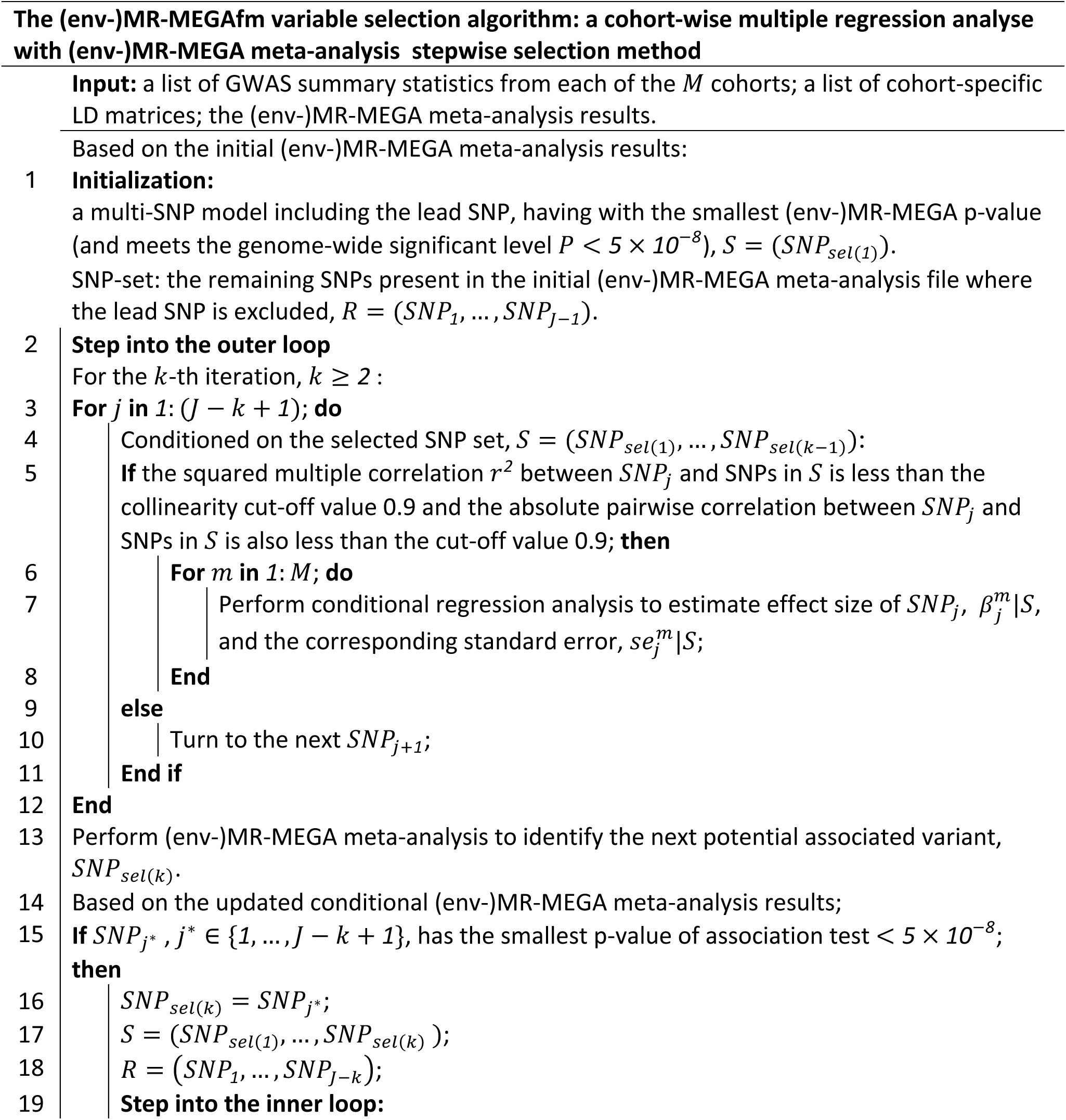
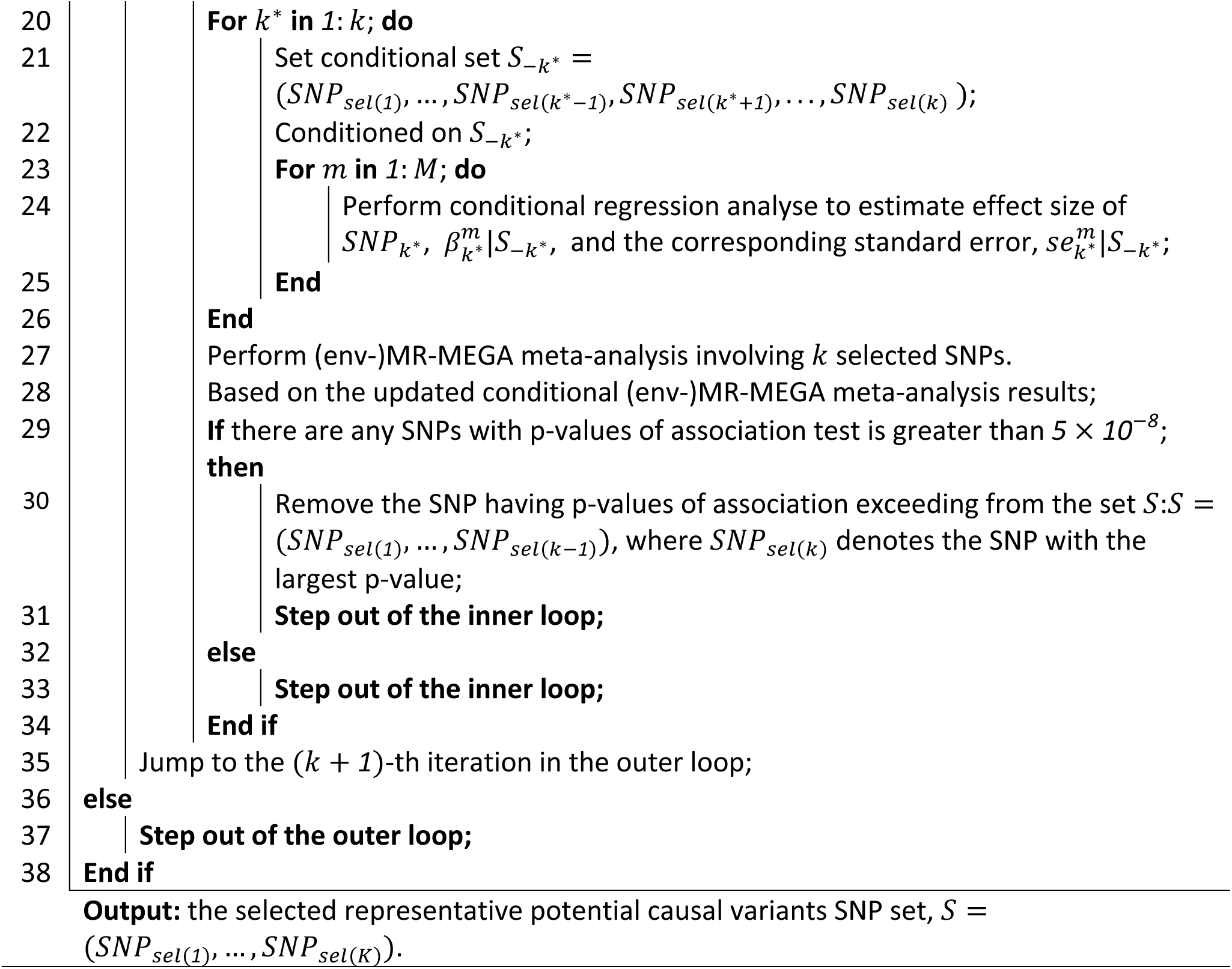

## Simulation design

To assess the performance of MR-MEGAfm and env-MR-MEGAfm, we conducted extensive simulations of multiple cohorts by varying allelic effects and different heterogeneity scenarios. All simulations were carried out under a realistic scenario that mimics the MAF in a region harbouring *APOE* (chr19: 44861224-45861224). For this region, we generated five continental African populations of 20,000 individuals based on 1000 Genomes Phase 3 reference panels^30^ using Hapgen2^31^: Esan in Nigeria (ESN), Gambian in Western Division (GWD), Luhya in Webuye (LWK), Mende in Sierra Leone (MSL), Yoruba in Ibadan (YRI) (Supplementary Figure1). In the subsequent simulation settings, we split the genotype data in each population into male and female cohorts with a total sample size of 10,000. The number of replications was set to 300. For each replication, across all populations, we randomly chose two causal variants from the genetic variants with MAF > 0.05 in each population, and for which the pair of causal variants satisfied *r^2^* < *0.6* within each population group. To simulate sex-stratified samples, traits were generated in a random sample of 5,000 individuals from each sample of 10,000 individuals.

Considering that allelic heterogeneity between populations is often correlated with environmental exposures beyond ancestry, we simulated the quantitative traits using the model of smoker-status association with the trait: *y*_*i*_ = *e*_*i*_ × (*β_1_* × *x*_*i*_*_1_*+ *β_2_*× *x*_*i*_*_2_*) + *ε*_*i*_, where *x_*i*j_* denotes the number of alternate alleles of variant *j* for individual *i* (i.e. genotype score), *e*_*i*_ is the indicator for smoking status of individual *i*: *e*_*i*_ = *1*(smoker)/0(non-smoker) and *β_j_*, *j* = *1*,*2* refers to the effects of the two causal variants. To mimic a realistic smoking setting, female cohorts had much lower smoking proportions compared to male cohorts^47^ (Supplementary Table 1). Within each replication, all variants that are present in the cohort with MAF>0.01 were assessed for association with the trait by MR-MEGA meta-analysis^25^, using two axes of genetic variation, and by env-MR-MEGA meta-analysis^27^ with the same axes of genetic variation along with two environmental covariates: cohort-level sex indicators and smoking proportions. It is noted that the axes of genetic variation used in MR-MEGA/env-MR-MEGA were derived from a matrix of mean pairwise allele frequency differences between cohorts, based on the genome-wide variants with MAF>0.05 across all African cohorts from 1000 Genomes Phase 3 reference panels^30^, rather than from the variants present within the fine-mapping region used.

To evaluate the accuracy of MR-MEGAfm and env-MR-MEGAfm, we compared the false discovery rates (FDR), defined as FDR is defined as the mean proportion of variants having marginal posterior probability (MPP) above a certain threshold that are non-causal variants, and the power, defined as the mean proportion of causal variants having MPP above a certain threshold (e.g. 0.5, 0.9). For each replication, a CS99 for each selected variant is constructed by including variants in decreasing order of their MPP until the cumulative sum first passes above 0.99. For ease in comparisons, within each replication, all CS99 are merged into the union of CS99. The coverage is estimated by the proportion of simulation replications in which the union of CS99 contains both causal variants and resolution is measured as the number of variants in the CS99 union.

As multiple credible sets are constructed (one based on each selected variant, conditional on the other selected variants) each variant has a marginal posterior probability (MPP) of causality calculated conditional on the other selected variants. To assess prioritisation of the causal variants, we define MPP distribution as the minimum of MPP between both causal variants, and MPP distribution indicates the frequency with which both causal variants exceed a given threshold.

To assess the performances of the proposed methods over a range of scenarios for heterogeneity in allelic effects between diverse populations, we introduce three heterogeneity scenarios across the five continental African populations, parameterised in terms of *β_j_* in each population (Supplementary Table S2): the ancestral homogeneity, west-centralAfrica and non-ancestral Africa scenarios. Under the ancestral homogeneity scenario, the allelic effects of causal variants are homogeneous across all cohorts. Under the west-central Africa scenario, the causal effects are specific to the west-central African populations (i.e. ESN and YRI). Under the non-ancestral Africa scenario, the causal effects are specific to one cohort from each African region - LWK, ESN and GWD.

For each heterogeneity scenario where allelic heterogeneity is impacted by smoking status, we assessed the performance of MR-MEGAfm, env-MR-MEGAfm and SuSiEx^15^ method in terms of coverage/calibration, power, resolution and FDR. It is noted that, in our implementation of SuSiEx, we set LEVEL option as 0.99 to ensure consistency with CS99 of (env-)MR-MEGAfm, MINIMUM_PURITY option as 0.0001 and KEEP_AMBIGUOUS_SNPS option as TRUE, which avoids filtering out SNPs on purity and ambiguous SNPs.

### African population data application

We applied these methods to sex-stratified summary statistics GWAS of LDL-cholesterol in 19,589 participants from five diverse sub-Saharan African cohorts. These are Burkina Faso, Ghana, Kenya, and South Africa in the Africa Wits-INDEPTH partnership for Genomics studies (AWI-Gen) consortium^34,35^, South African Zulu cohorts: the Durban Diabetes Case Control Study (DCC) and the Durban Diabetes Study (DDS)^36^ and Uganda: Uganda Genome Resource^37–39^. Detailed descriptions of these study cohorts have been published, including the ethical consent, community engagement, genotyping methods, quality control, imputations^36^, and the GWAS performed for each cohort. Briefly, we aggregated data from African cohorts: AWI-Gen, South African Zulu, and the Uganda Genome Resource.

The AWI-Gen cohort included 10,898 participants from three sub-Saharan African centres: South Africa (5,265 participants), East Africa (1,766 participants), and West Africa (3,867 participants). The African Genome Resources reference panel was used for the imputation, resulting in 13.98 million high-quality SNPs after stringent filtering. The South African Zulu cohort comprised 2,707 individuals from the Durban Diabetes Study and Case Control Study. After quality control, 16,559,897 variants and 2,572 participants were retained. The Uganda Genome Resource cohort comprised 7,000 individuals, resulting in 6,119 individuals for analysis. The final dataset contained 15,783,409 SNPs after excluding those with a minor allele frequency below 1%. Residuals from LDL-C regression on age and principal components were inverse rank normalized, and genome-wide association analysis was performed using sex - and cohort-stratified linear mixed models in GCTA^32^.

The summary statistics obtained from the MR-MEGA^25^ and env-MR-MEGA^27^ meta-analysis of the twelve sex-stratified GWAS of LDL-cholesterol were used to fine-map signals in two previously identified regions^27^. Briefly, to generate axes of genetic variation, we filtered out SNPs with MAF below 5% within each cohort and removed ambiguous SNPs (i.e. A/T, G/C). We performed meta-analysis at non-ambiguous SNPs with MAF>0.01 across all cohorts using MR-MEGA and env-MR-MEGA methods; env-MR-MEGA included the urban status proportions to account for environmental factors that could introduce heterogeneity of allelic effects. We stratified by sex for a meta-analysis involving 12 cohorts, generating p-values for LDL-cholesterol association and heterogeneity due to ancestry and environmental variables.

For the genomic regions, we focused on the 1Mb regions centred on the two genome-wide significant loci on chromosome 1 (*PCSK9*: rs28362286) and chrmosome 19 (*APOE:* rs7412). We then extracted the summary-level data and the true cohort-level for each region as inputs for the (env-)MR-MEGAfm. It is noted that, in previous analyses^27^ rs7412 in *APOE* showed significant evidence of heterogeneity due to genetic ancestry and environment (proportion of participants residing in urban areas) whereas rs28362286 in *PCSK9* had evidence of heterogeneity due to urban status only. Additionally, we have also applied env-MR-MEGA using continuous environmental covariates (BMI)^27^. We further confirmed that the genomic region boundaries which are recommended by SABER^40^ align with the predefined genomic regions.

We constructed each cohort-level LD matrix (not sex-stratified) for fine-mapping using PLINK v1.9b^48,49^. The number of variants used to generate LD matrices for the AWI-Gen cohorts was identical, as the three African sub-regions were processed and genotyped together, including pre-and post-imputation quality control procedures, retaining only common SNPs across the study sites and countries^50^. Specifically, for the rs28362286 region, the LD matrix for the region centered on rs28362286 included 7,301 variants in the Eastern, Southern, and Western AWI-Gen cohorts, 15,221 variants in the South African Zulu cohort, and 8,323 variants in the Uganda cohort. For the rs7412 region, the LD matrix for the region centered on rs7412 included 6,885 variants in the Eastern, Southern, and Western AWI-Gen cohorts, 16,516 variants in the South African Zulu cohort, and 8,884 variants in the Uganda cohort.

## Notes

### Competing Interest Statement

The authors have declared no competing interest.

