## Supplementary material for "Multi-ancestry fine-mapping accounting for ancestral and environmental heterogeneity improves resolution and interpretation"

**Supplementary Figure 1. Axes of genetic variation separating five continental African ancestry populations.** The first two axes of genetic variation from multi-dimensional scaling of the Euclidean distance matrix between the ten sex-stratified cohorts are sufficient to separate population groups from different regions of Africa: east Africa: Luhya in Webuye, Kenya (LWK), west-central Africa: Esan in Nigeria (ESN) and Yoruba in Ibadan, Nigeria (YRI) and west Africa: Gambian in Western Division, The Gambia (GWD) and Mende in Sierra Leone (MSL).

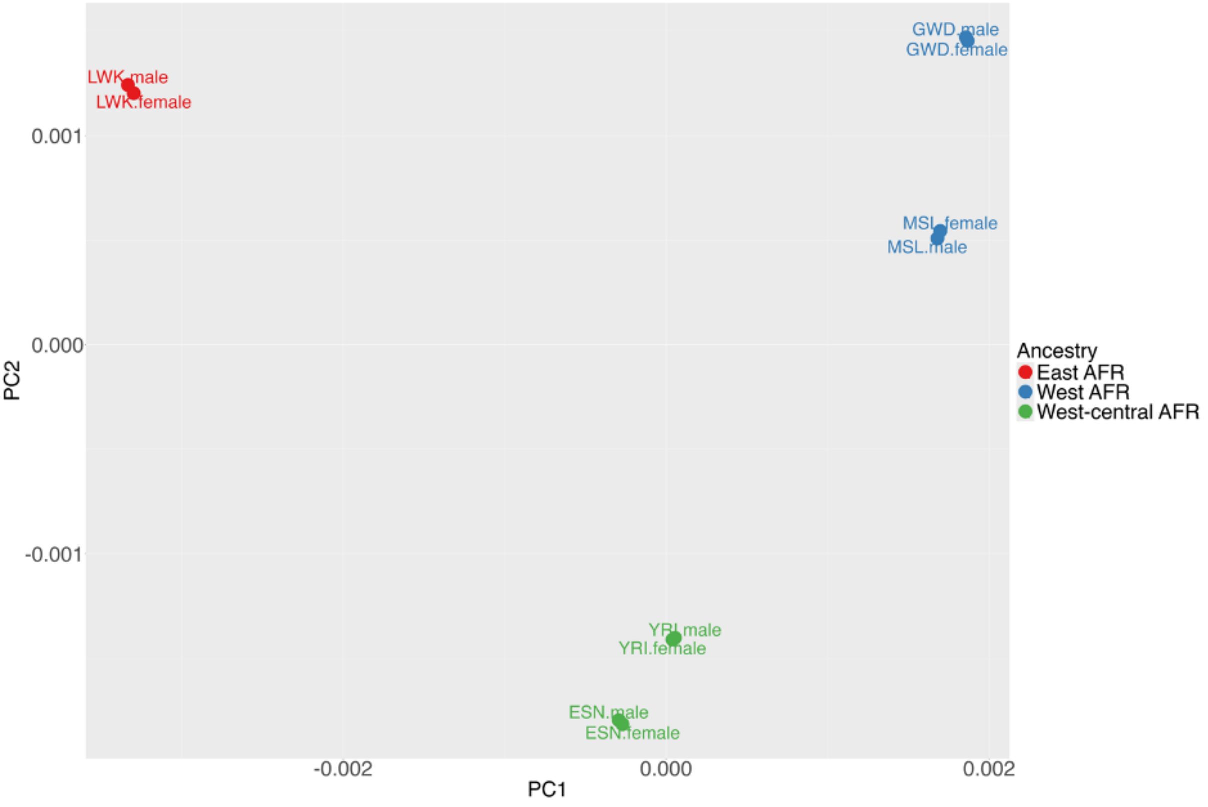

**Supplementary Figure 2. Under the homogeneous scenario where allelic heterogeneity is uncorrelated with environment, env-MR-MEGAfm and MR-MEGAfm had well-calibrated coverage at 99%. Meanwhile, all fine-mapping methods shared comparable performance in prioritisation of both causal variants. (a). Coverage from the homogeneous scenario without environmental effects.** Coverage is measured as the probability that both causal variants are captured by the CS99 union estimated over 300 replications. Data are presented as the proportion of replications in which the CS99 union contains all causal variants SEM, where SEM is the standard proportion error bound of a 95% confidence interval based on 300 replications. (**b). Prioritisation Assessment.** The violin plots show MPP distribution of the three fine-mapping methods, and the median is given by the centre line, upper and lower quartiles are the box limits, and whiskers are at most interquartile range. This suggests that when the quantitative traits are not affected by both ancestry and environmental exposures, env-MR-MEGAfm has comparable performance of prioritisation of both causal variants compared to MR-MEGAfm. Additionally, there is no significant difference in prioritisation between the proposed fine-mapping methods, MR-MEGAfm/env-MR-MEGAfm and SuSiEx.

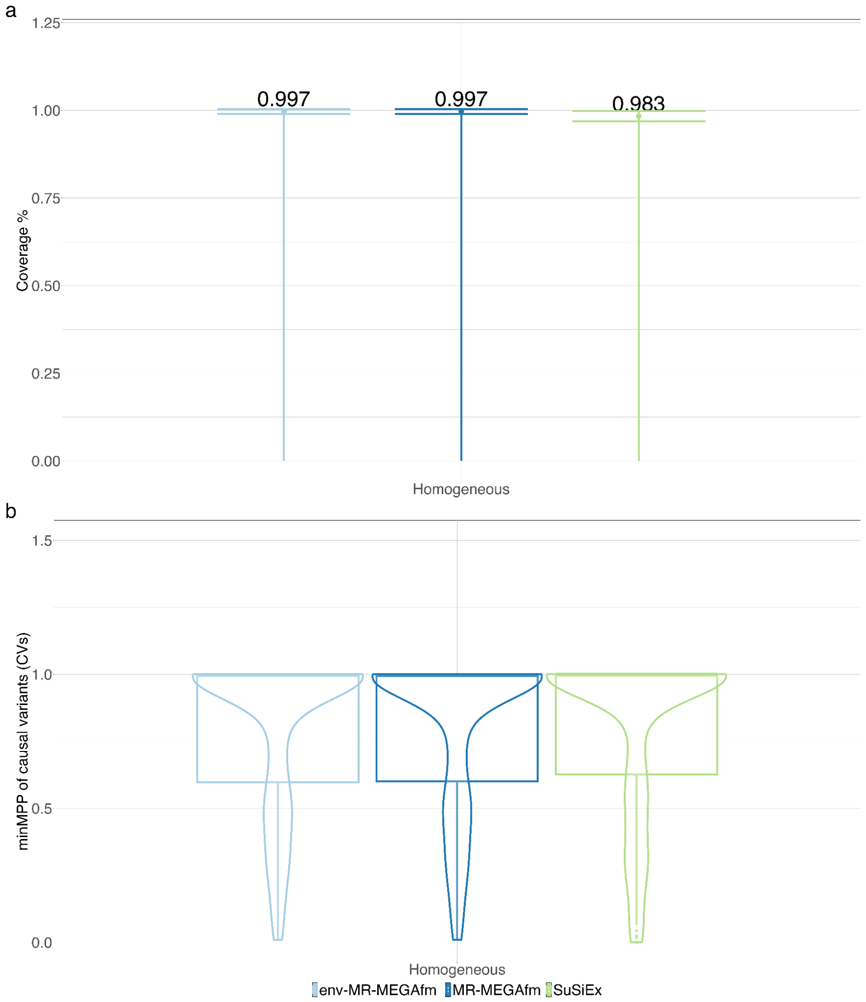

**Supplementary Figure 3. Under the homogeneous scenario without environmental effects, across 300 replications, the sizes of CS99 union from env-MR-MEGAfm were comparable with that from MR-MEGAfm. However, compared to env-MR-MEGAfm, the sizes of CS99 union from SuSiEx were substantially larger than those from env-MR-MGAfm. a. Resolution Assessment**. Distribution of CS99 union sizes from env-MR-MEGAfm, MR-MEGAfm and SuSiEx; the median is given by the centre line, upper and lower quartiles are the box limits, and whiskers are at most  interquartile range. **b**. **Prioritisation Assessment.** The scatter plot for comparison of CS99 union between env-MR-MEGAfm and SuSiEx. This indicates that most of CS99 union sizes from env-MR-MEGAfm are significantly smaller than those from SuSiEx.

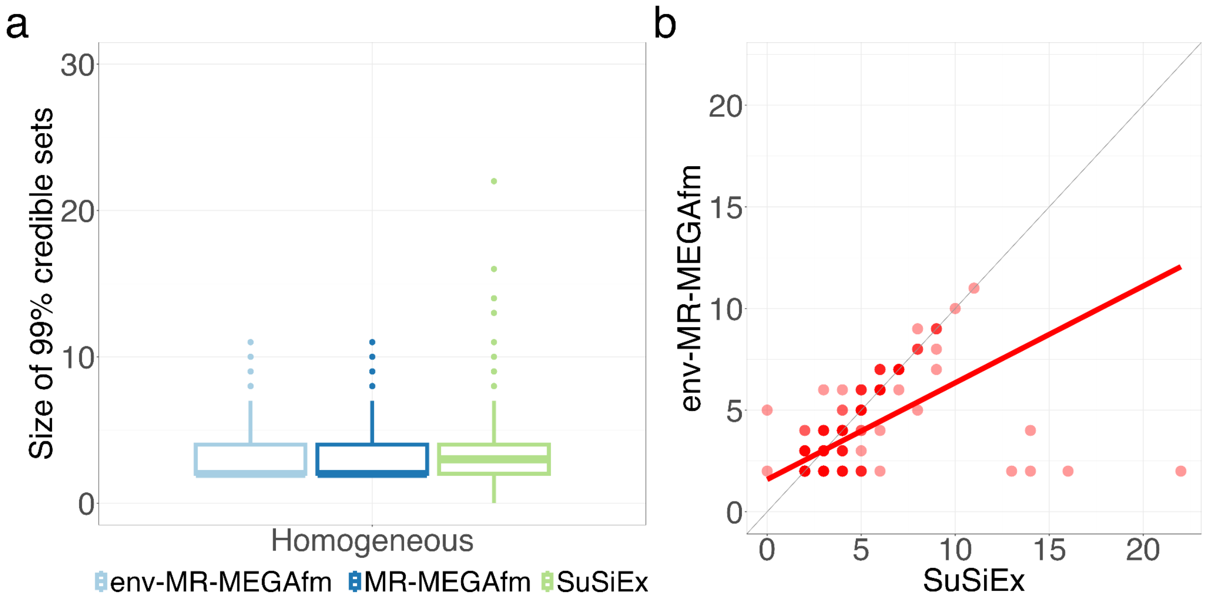

**Supplementary Figure 4. Across varied allelic effects settings, under the ancestrally homogeneous scenario with varying environmental effects, env-MR-MEGAfm always had higher gains in resolution and prioritisation over MR-MEGAfm. Additionally, as the allelic effects increased, both methods had improved resolution and prioritisation. Compared to the proposed fine-mapping methods, SuSiEx had comparable performance in resolution and prioritisation compared with env-MR-MEGAfm**. Within each setting, there are 300 replications; any pair of causal variants has the same allelic effects. **a. Resolution Assessment**. Distribution of CS99 union sizes from env-MR-MEGAfm ,MR-MEGAfm and SuSiEx; the median is given by the centre line, upper and lower quartiles are the box limits, and whiskers are at most interquartile range. This indicated that env-MR-MEGAfm and SuSiEx had better resolution than MR-MEGAfm regardless of causal effects. **b Prioritisation Assessment.** The boxplots show MPP distribution of the three fine-mapping methods, and the median is given by the centre line, upper and lower quartiles are the box limits, and whiskers are at most interquartile range. This suggested gains in prioritisation of causal variants by env-MR-MEGAfm and SuSiEx, compared to MR-MEGAfm, at large allelic effect sizes.

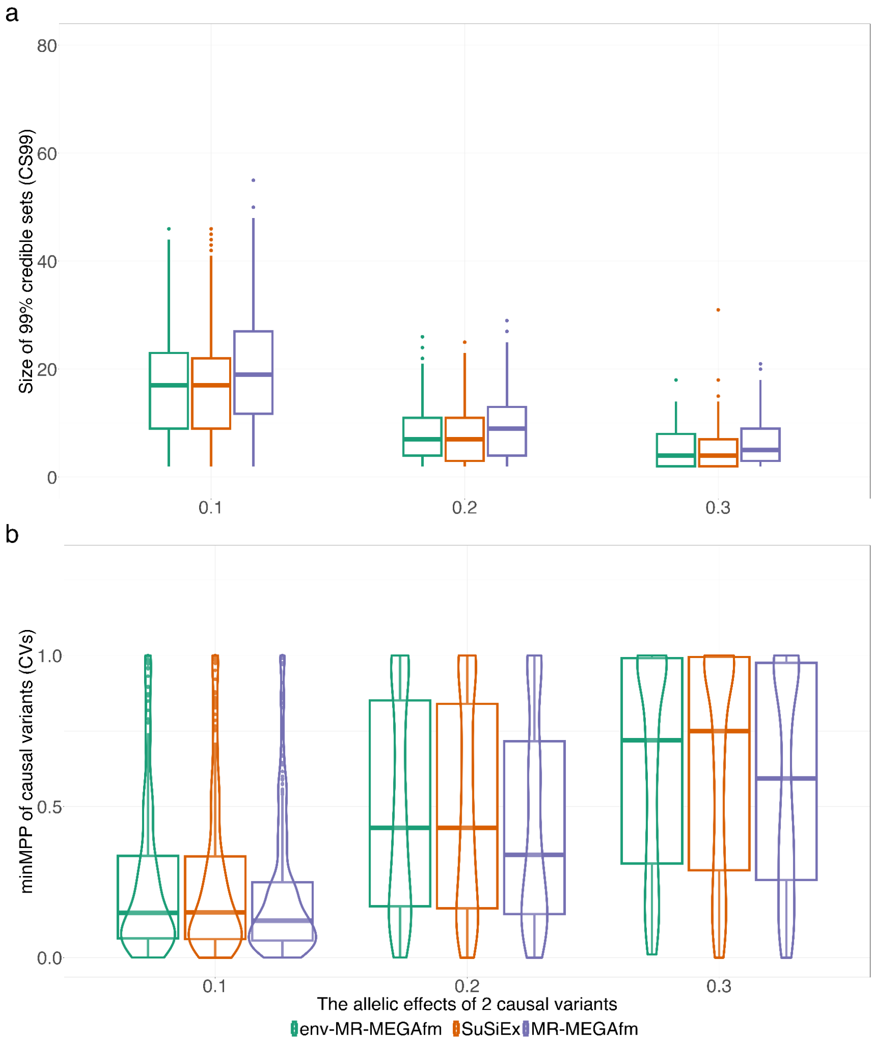

**Supplementary Figure 5. Under the ancestrally homogeneous scenario where heterogeneity of allelic effects is correlated with environment only, env-MR-MEGAfm had comparable CS99 union sizes with SuSiEx when causal effects ranging from 0.1 to 0.2, but notably produced smaller CS99 union sizes when causal effects were 0.3.** The CS99 union sizes are constructed by counting the number of variants included in all CS99 from env-MR-MEGAfm and SuSiEx.

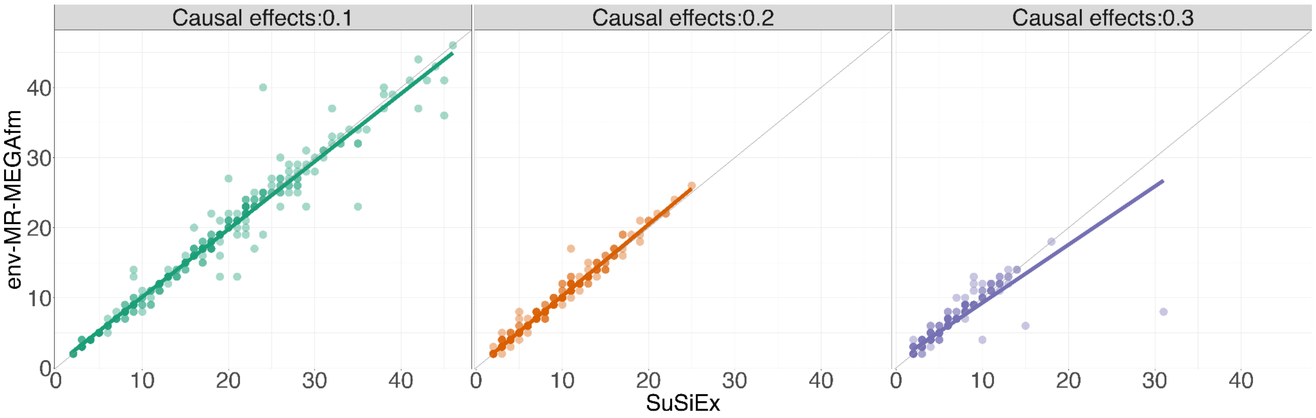

**Supplementary Figure 6. Across different heterogeneity scenarios, both env-MR-MEGAfm and SuSiEx had comparable resolution performance, while the resolution of env-MR-MEGAfm was significantly higher than that of MR-MEGAfm.** The scatter plot shows a comparison of CS99 union sizes between env-MR-MEGAfm and the other two fine-mapping methods: MR-MEGAfm and SuSiEx. This indicates that env-MR-MEGAfm and SuSiEx consistently have better resolution than MR-MEGAfm regardless of heterogeneity scenarios.

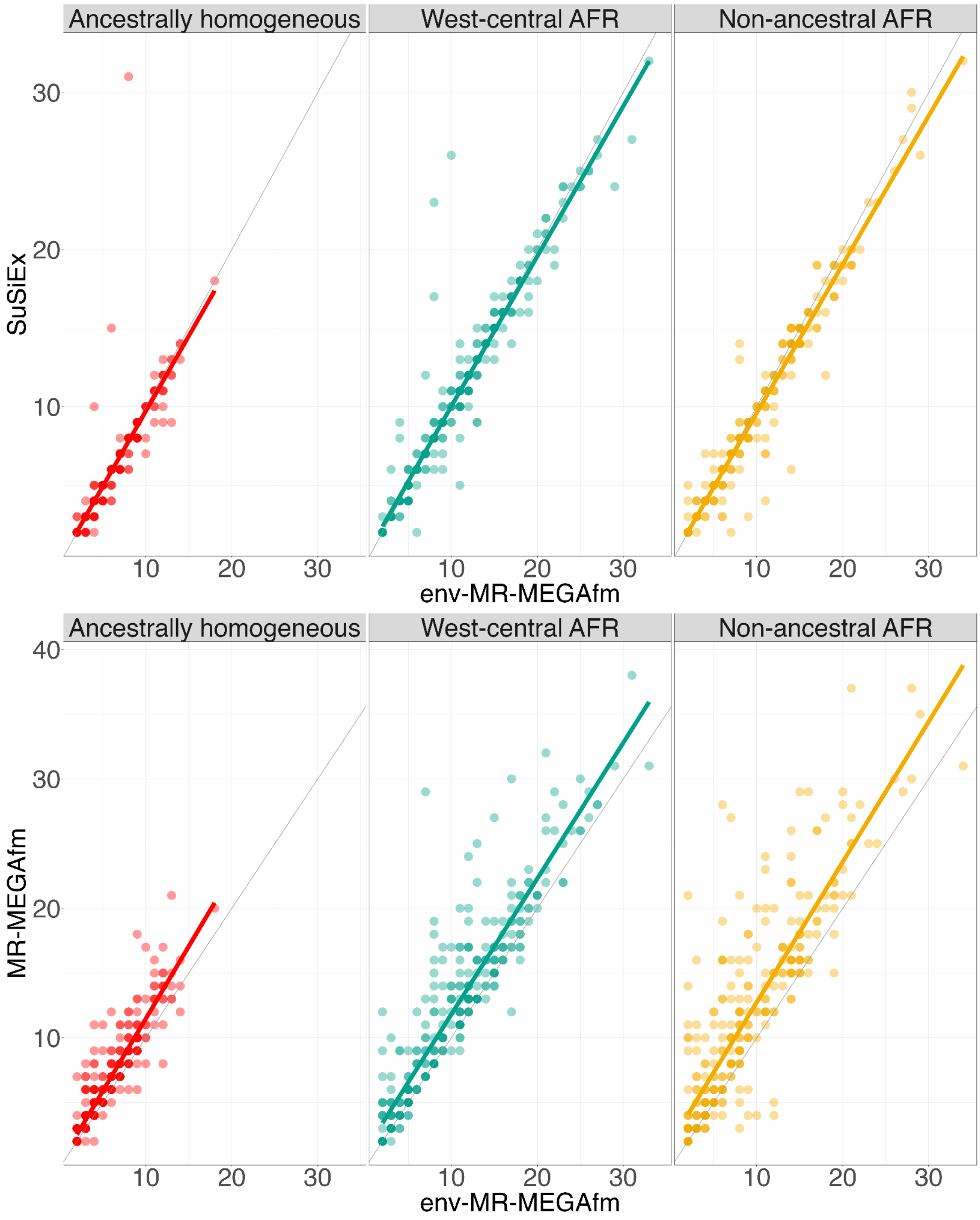

**Supplementary Figure 7. Across different heterogeneity scenarios, env-MR-MEGAfm had comparable performance in prioritisation of both causal variants with SuSiEx.** The X-axis refers to MPP of causal variants from SuSiEx and the Y-axis refers to MPP of causal variants from env-MR-MEGAfm. The red line refers to the fitted line. This indicates that, across 300 replications, most of MPP from env-MR-MEGAfm are nearly equivalent with those from SuSiEx.

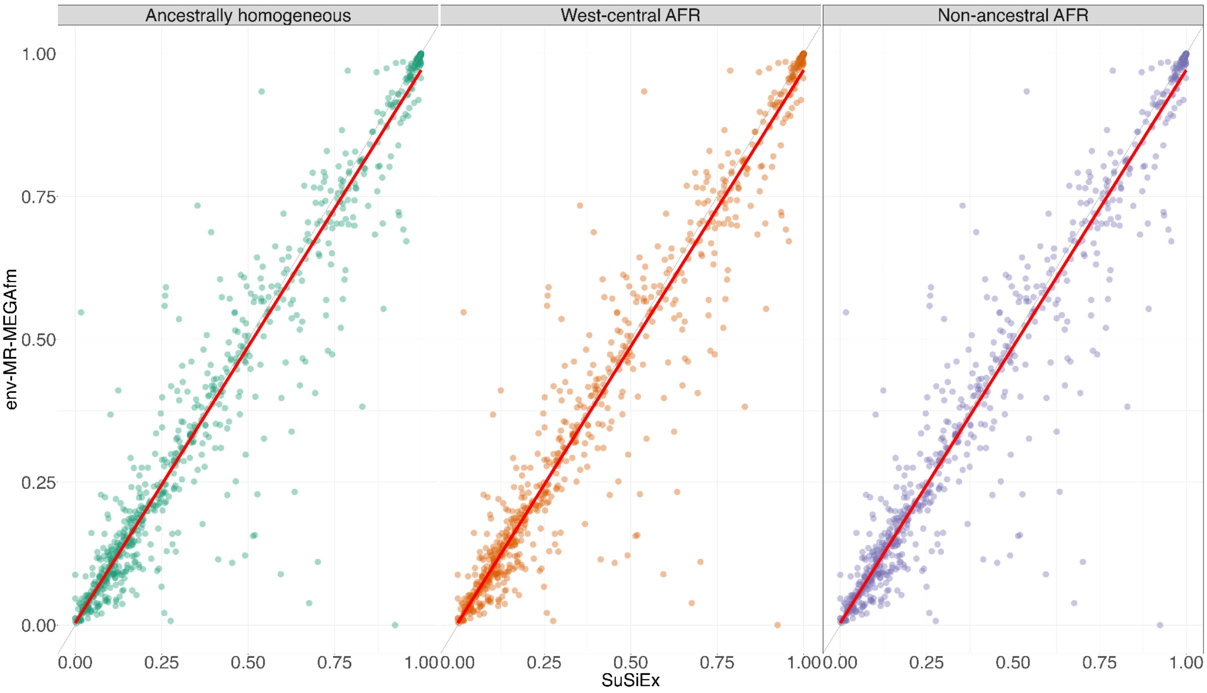

**Supplementary Figure 8. Under the ancestrally homogeneous scenario, utilization of the combined LD in env-MR-MEGAfm, MR-MEGAfm and SuSiEx gives a notable reduction in resolution and prioritisation compared to the true cohort-level LD. a Distribution of CS99 union sizes from env-MR-MEGAfm and MR-MEGAfm**; the median is given by the centre line, upper and lower quartiles are the box limits, and whiskers are at most the interquartile range. **b Prioritisation Assessment.** The three boxplots show MPP distribution of the three fine-mapping methods, and the median is given by the centre line, upper and lower quartiles are the box limits, and whiskers are at most interquartile range. This suggests that, regardless of the fine-mapping methods, the utilization of the true cohort-level LD showed significantly improved prioritization than that of the combined LD.

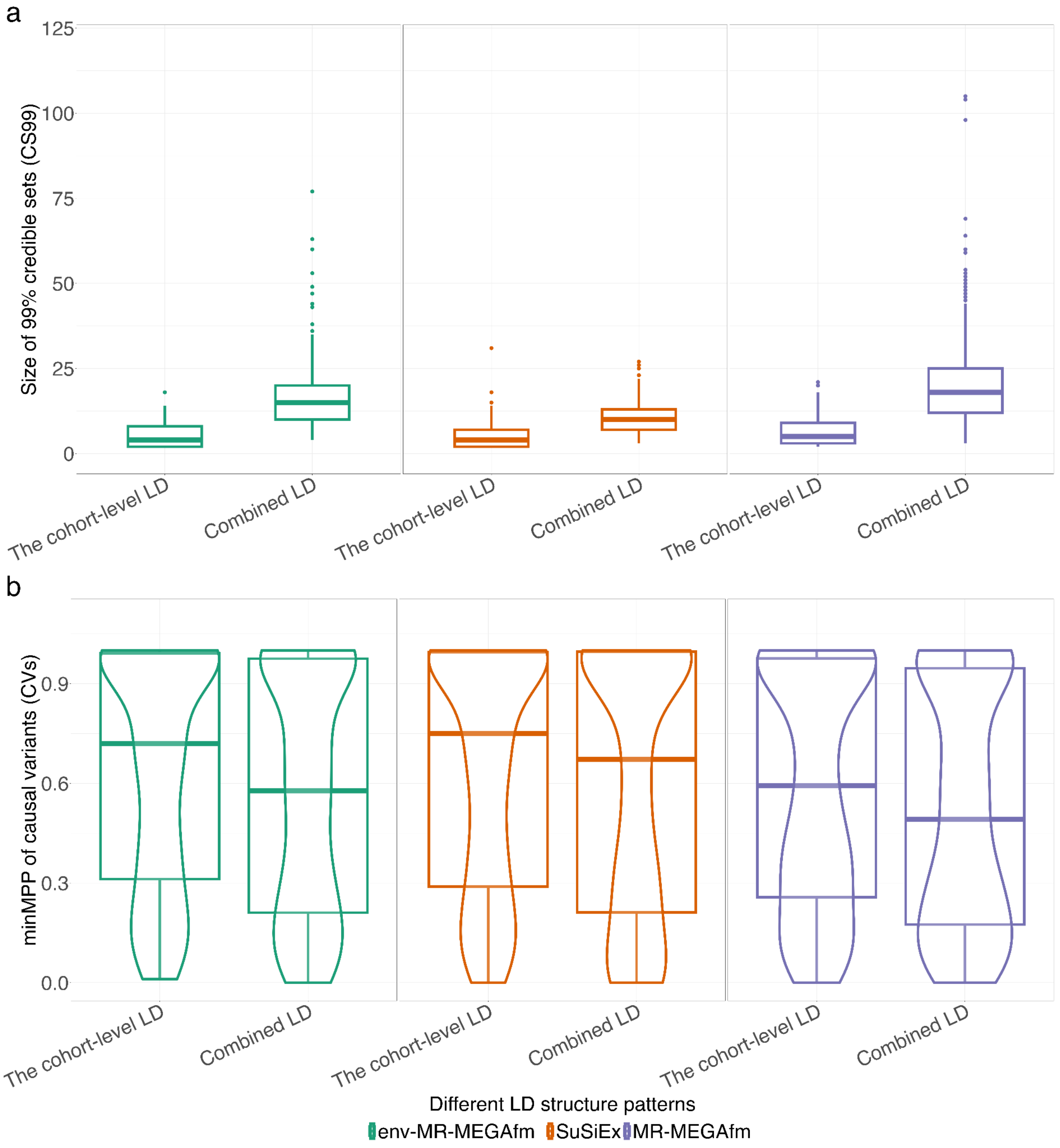

**Supplementary Table 1. Simulation settings for smoking proportions in female and male cohorts in which five populations from the continental African region were stratified by sex.** To mimic the realistic setting, female cohorts across all populations have less smokers than male cohorts.

| Populations | | | Smoke proportions (female/male) |
|---|---|---|---|
| Code | Description | Region | Same direction |
| LWK | Luhya in Webuye, Kenya | East Africa | 0.243/0.81 |
| ESN | Esan in Nigeria | West-central Africa | 0.215/0.836 |
| YRI | Yoruba in Ibadan, Nigeria | West-central Africa | 0.261/0.888 |
| GWD | Gambian in Western Division, The Gambia | West Africa | 0.192/0.676 |
| MSL | Mende in Sierra Leone | West Africa | 0.256/0.655 |

**Supplementary Table 2. Both env-MR-MEGAfm and MR-MEGAfm using the true cohort-level LD were well-calibrated regardless of causal effects and heterogeneity scenarios. As expected, both methods using the combined LD showed minor reduction in coverage. Additionally, SuSiEx showed comparable performance of calibration compared to the proposed fine-mapping methods under different causal effect settings and varied heterogeneity scenarios, while SuSiEx showed lower coverage than env-MR-MEGAfm and MR-MEGAfm.** Coverage is measured as the probability that both causal variants are captured by the CS99 union estimated over 300 replications. The values in the parentheses bracket refers to the 95% confidence interval for the coverage estimate - the proportion of replications in which the CS99 union contains both causal variants ± SEM, where SEM is the standard proportion error bound of a 95% confidence interval based on 300 replications.

<table>
<tr><td colspan="4">(a) Varying causal effects $(\beta_1, \beta_2)$ under the ancestrally homogeneous scenario</td></tr>
<tr><td>$(\beta_1, \beta_2)$</td><td>(0.1, 0.1)</td><td>(0.2, 0.2)</td><td>(0.3, 0.3)</td></tr>
<tr><td>MR-MEGAfm</td><td>0.987<br>(0.979,1.000)</td><td>0.996<br>(0.990,1.003)</td><td>0.993<br>(0.984,1.003)</td></tr>
<tr><td>env-MR-MEGAfm</td><td>0.990<br>(0.979,1.001)</td><td>0.996<br>(0.990,1.003)</td><td>1.000<br>(1.000,1.000)</td></tr>
<tr><td>SuSiEx</td><td>0.990<br>(0.979,1.001)</td><td>0.996<br>(0.990,1.003)</td><td>0.996<br>(0.990,1.003)</td></tr>
<tr><td colspan="4">(b) Varying ancestral heterogeneity scenarios under the causal effect setting where $(\beta_1, \beta_2)$=(0.3, 0.3)</td></tr>
<tr><td></td><td>Ancestrally homogeneous</td><td>West-central Africa</td><td>Non-ancestral Africa</td></tr>
<tr><td>MR-MEGAfm</td><td>0.993<br>(0.984,1.003)</td><td>0.987<br>(0.974,1.000)</td><td>0.957<br>(0.934,0.980)</td></tr>
<tr><td>env-MR-MEGAfm</td><td>1.000<br>(1.000,1.000)</td><td>1.000<br>(1.000,1.000)</td><td>0.997<br>(0.990,1.003)</td></tr>
<tr><td>SuSiEx</td><td>0.996<br>(0.990,1.003)</td><td>0.993<br>(0.984,1.003)</td><td>0.993<br>(0.984,1.003)</td></tr>
<tr><td colspan="4">(c) Different LD patterns under the ancestrally homogeneous scenario</td></tr>
<tr><td></td><td>The true cohort-level LD</td><td colspan="2">Combined LD</td></tr>
</table>

| | | |
|---|---|---|
| MR-MEGAfm | 0.993<br>(0.984,1.003) | 0.987<br>(0.974,1.000) |
| env-MR-MEGAfm | 1.000<br>(1.000,1.000) | 0.983<br>(0.969,0.998) |
| SuSiEx | 0.996<br>(0.990,1.003) | 0.930<br>(0.901,0.959) |

**Supplementary Table 3. Across varied causal effect settings, under the ancestrally homogeneous scenario where heterogeneity of allelic effects is correlated with environment only, env-MR-MEGAfm always had comparable power with SuSiEx, while it demonstrated higher power than MR-MEGAfm.** As the causal effects increased, power yielded by the three fine-mapping methods became higher while FDR kept comparably stable, regardless of threshold. FDR is defined as the mean proportion of non-causal variants having MPP above a certain threshold, and power is measured as the mean proportion of causal variants having MPP above a certain threshold. Estimates are based on 300 simulation replications.

| Causal effects ($\beta_1, \beta_2$) | 0.3 | | 0.2 | | 0.1 | |
|---|---|---|---|---|---|---|
| Threshold (0.5) | FDR | Power | FDR | Power | FDR | Power |
| MR-MEGAfm | 0.088 | 0.740 | 0.102 | 0.645 | 0.123 | 0.377 |
| env-MR-MEGAfm | 0.072 | 0.795 | 0.107 | 0.673 | 0.113 | 0.433 |
| SuSiEx | 0.082 | 0.800 | 0.108 | 0.683 | 0.121 | 0.437 |
| Threshold (0.9) | | | | | | |
| MR-MEGAfm | 0.010 | 0.583 | 0.003 | 0.442 | 0.000 | 0.192 |
| env-MR-MEGAfm | 0.003 | 0.635 | 0.003 | 0.498 | 0.007 | 0.238 |
| SuSiEx | 0.018 | 0.647 | 0.005 | 0.503 | 0.005 | 0.240 |

**Supplementary Table 4. Heterogeneity scenarios parameterized in terms of $\beta$ in each continental African population. For the five continental African populations, west-central Africa (ESN,YRI), west Africa (GWD,MSL), and east Africa (LWK) were collected from Phase 3 of the 1000 Genomes project.**

| Populations | | Ancestrally homogeneous | West-central AFR | Non-ancestral AFR |
|---|---|---|---|---|
| Code | Region | | | |
| LWK | East Africa | $\beta$ | 0 | $\beta$ |
| ESN | West-central Africa | $\beta$ | $\beta$ | $\beta$ |
| YRI | West-central Africa | $\beta$ | $\beta$ | 0 |
| GWD | West Africa | $\beta$ | 0 | $\beta$ |
| MSL | West Africa | $\beta$ | 0 | 0 |

**Supplementary Table 5. Two sample size settings across the ten sex-stratified African population groups.** In Sample size setting 1, the sample sizes of all African cohorts are consistent with the real sample sizes of specified African regions. For example, the east Africa population group has around 2,500 females and 3,500 males, which is assigned to LWK sample size. In Sample size setting 2, we set smaller sample sizes across ten sex-stratified cohorts.

| Population groups | Sex | Sample size setting 1 | Sample size setting 2 |
|---|---|---|---|
| ESN | Female | 2,000 | 1,000 |
| | Male | 2,000 | 1,000 |
| GWD | Female | 2,000 | 1,000 |
| | Male | 2,000 | 1,000 |
| LWK | Female | 2,500 | 1,000 |
| | Male | 3,500 | 1,000 |
| MSL | Female | 2,000 | 1,000 |
| | Male | 2,000 | 1,000 |
| YRI | Female | 1,000 | 1,000 |
| | Male | 1,000 | 1,000 |

**Supplementary Table 6. Under west-central Africa scenario where causal effects were specific to west-central Africa population groups (ESN, YRI), regardless of sample size settings (sample size per cohort ≤3,500), all fine-mapping methods were well-calibrated at coverage of 99%, with the upper limit exceeding 0.95. Under different thresholds, all fine-mapping methods had comparable FDR and power. Specifically, env-MR-MEGAfm and SuSiEx had comparable results of FDR, power and coverage. Additionally, they produced higher power and better calibration at coverage than MR-MEGAfm.** Estimates are based on 300 simulation replications.

| Scenarios | Sample size setting 1 | | | Sample size setting 2 | | |
|---|---|---|---|---|---|---|
| Threshold (0.5) | FDR | Power | Coverage | FDR | Power | Coverage |
| MR-MEGAfm | 0.100 | 0.292 | 0.960 (0.938,0.982) | 0.118 | 0.253 | 0.957 (0.934,0.980) |
| env-MR-MEGAfm | 0.153 | 0.325 | 0.973 (0.955,0.992) | 0.107 | 0.283 | 0.967 (0.946,0.987) |
| SuSiEx | 0.137 | 0.358 | 0.993 (0.984,1.003) | 0.113 | 0.297 | 0.970 (0.957,0.989) |
| Threshold (0.9) | | | | | | |
| MR-MEGAfm | 0.010 | 0.153 | | 0.013 | 0.123 | |
| env-MR-MEGAfm | 0.010 | 0.193 | | 0.010 | 0.160 | |
| SuSiEx | 0.002 | 0.218 | | 0.007 | 0.167 | |

**Supplementary Table 7. Compared to the results under the ancestrally homogeneous scenario with the true cohort-level LD patterns as inputs, env-MR-MEGAfm MR-MEGAfm and SuSiEx implemented with the combined LD provided slightly higher FDR and lower power, regardless of threshold.** FDR is defined as the mean proportion of variants having marginal posterior probability (MPP) above a certain threshold that are non-causal variants, and power is measured as the mean proportion of causal variants having MPP above a certain threshold. Estimates are based on 300 simulation replications.

| LD pattern | The true cohort-level LD | | Combined LD | |
|---|---|---|---|---|
| Threshold (*0.5*) | FDR | Power | FDR | Power |
| MR-MEGAfm | 0.088 | 0.740 | 0.659 | 0.692 |
| env-MR-MEGAfm | 0.072 | 0.795 | 0.654 | 0.745 |
| SuSiEx | 0.082 | 0.800 | 0.709 | 0.760 |
| Threshold (*0.9*) | | | | |
| MR-MEGAfm | 0.010 | 0.583 | 0.587 | 0.538 |
| env-MR-MEGAfm | 0.003 | 0.635 | 0.603 | 0.583 |
| SuSiEx | 0.018 | 0.647 | 0.724 | 0.645 |
